# Intrinsic coordination of dynamic molecular signatures shape the human prefrontal cortex

**DOI:** 10.64898/2026.05.13.724991

**Authors:** Patricia R. Nano, Daniel C. Jaklic, Vanna Giang, Jose A. Soto, Jessenya Mil, Sean Wang, Antoni Martija, Brittney Wick, Maximilian Haeussler, Aparna Bhaduri

## Abstract

The cerebral cortex drives human cognition through the coordinated activity of discrete cortical areas, each harboring specialized molecular, structural and functional characteristics. Central to this organization is the prefrontal cortex (PFC), a hub for executive function that displays disproportionate expansion in humans and selective vulnerability to neurodevelopmental disorders. Previous work has identified a collection of PFC-enriched marker genes with dynamic expression trajectories, and re-analysis of these datasets converge these markers into 18 distinct molecular signatures of spatiotemporal PFC identity. However, the intrinsic gene networks that coordinate these molecular signatures to shape the human PFC remains unclear. Through pooled CRISPR activation screens in human primary cortical tissues, we have evaluated the ability of PFC-enriched transcription factors to intrinsically pattern PFC molecular identity. Our screens identify novel roles for the neurogenesis regulator, YBX1, in the activation of human PFC fate. In parallel screens and knock-down experiments in human cortical organoids, we define how YBX1 acts in concert with other PFC determinants to activate molecular signatures of PFC identity. Our findings support a model in which PFC patterning is orchestrated by cohorts of intrinsic determinants that initiate, potentiate, and modulate PFC gene signatures, conferring robustness to the development of the human PFC.

## Introduction

The prefrontal cortex (PFC) is a well-established hub of both our uniquely human cognitive capacities and vulnerability to neurological disease. Compared to other functionally specialized areas within the cortex, the PFC has unique synaptic and dendritic properties^1^, protracted maturation^2^, and ultimately, occupies the most space^3^. These distinctions are intensified in the human, which contains a massively expanded PFC relative to that of the mouse and even non-human primates^3^. While mouse models have laid the foundation for how the PFC can be patterned during cortical development^4–6^, understanding how the PFC drives the human ability to sense, feel, and make decisions requires investigating this cortical area in a human context. Motivated by these open questions, recent efforts to profile the human brain have often centered on the PFC^7–9^, resulting in an extensive catalog of PFC-enriched genes in development^7,8^ and disease^10–12^. How these molecular programs are orchestrated to specify and maintain PFC identity remains poorly understood.

The emergence of cortical areas, including the PFC, begins with rostrocaudal morphogen gradients such as FGF8 that drive initial patterning^13^. Competing models have long debated whether cortical arealization is driven by intrinsic transcription factors and signaling gradients present in the early cortical sheet (protomap model), or extrinsically, by thalamic and other sensory inputs that shape areal identity later in development (protocortex model). Exploration of these opposing forces in human cortical development has largely determined that consistent with the protomap, early intrinsic signals establish the rostrocaudal poles of the cortex while the intervening sensory regions may require extrinsic refinement^14,15^. These conclusions have primarily emerged from the transcriptomic profiling of primary samples of the developing human cortex^14–16^, with key experimental verification in mouse models linking signaling cascades to persistent PFC marker genes including *CBLN2* and *MEIS2*^17^.

While these cortical “arealization” processes have been examined at the level of bulk tissue^16^and single cells14,15, the congruence of single-cell to bulk transcriptomic profiles is generally low. Indeed, single-cell transcriptomic profiling has revealed the presence of more dynamic PFC marker genes, which are transiently enriched in the PFC at specific cell types or developmental stages^14,15^. We have previously hypothesized that these spatiotemporal dynamics underlie a domino effect, whereby sequential, PFC-enriched gene expression programs cascade across stages of peak neurogenesis^14^. However, these dynamic expression patterns have yet to be translated into tractable molecular readouts of PFC fate, limiting our ability to interrogate how they are regulated.

Efforts to progress from a descriptive to a mechanistic understanding of PFC development have been further constrained by the limited availability of human primary cortical tissue (HPCT). Human cortical organoids (hCOs) have emerged as a complementary and more accessible *in vitro* model, recapitulating key cell types and developmental transitions of the human cortex^18–20^. Cortical organoid cells express known areal marker genes, albeit in a stochastic and spatially disorganized manner^21^. Organoids have also been used to explore how extrinsic morphogens^22,23^ as well as key signaling molecules such as retinoic acid^17^ can promote PFC transcriptional identity. These prior mechanistic studies have highlighted how extrinsic cues can activate PFC-enriched gene programs; in contrast, the ability of intrinsic determinants to coordinate the dynamic gene expression patterns that underlie PFC fate remains largely untested.

Here, we employ network co-expression methods and pooled CRISPR activation screening to evaluate the dynamic molecular landscape of PFC identity. We implemented network co-expression methods^24^ to establish 18 cell type- and stage-specific PFC gene signatures, generating key metrics of PFC development. To identify the intrinsic regulators of these PFC signatures, we conducted pooled CRISPR activation screening of 35 PFC-enriched transcription factors (TF) in HPCT. The resulting perturbational dataset demonstrates the ability of intrinsic TFs to activate at least 15 of these 18 PFC signatures, which were primarily associated with the regulation of neurogenesis. We then conducted a parallel CRISPR activation screen in hCOs, benchmarking these findings to our HPCT screen to systematically isolate the features of PFC gene expression that are preserved and testable in hCOs. Emerging in both of these contexts is a novel role in human PFC specification for YBX1, a screen target with well-established roles in neurogenesis^25,26^. With complementary YBX1 knock-down experiments in hCOs, our data reveal that this individual TF is sufficient to activate a subset of PFC signatures, yet required for the maintenance of nearly all PFC signatures; and these regulatory functions are transduced and reinforced by cohorts of PFC-enriched TFs. By generating a perturbational dataset in HPCT and an analogous, benchmarked resource in hCO, these studies establish new, high-throughput approaches for extending the transcriptomic profiles of the developing human cortex into mechanisms of human cortical arealization.

## Results

### Derivation of spatiotemporally dynamic molecular signatures underlying human PFC development

We first sought to generate tractable, molecular readouts that can capture the dynamic gene expression patterns inherent in the developing PFC^14,15^. As a baseline, we evaluated how marker gene sets from previous bulk transcriptional studies dynamically change across frontal and occipital lobe dissections in single-cell datasets. These analyses used a bulk transcriptomic dataset of meticulously dissected laminar regions from the PFC and a number of other cortical areas^16^. We extracted a group of PFC-enriched genes, then cross-validated these with independently generated single-cell datasets of the developing PFC^14^, narrowing down this list to the genes that had the greatest effect (see Methods, **Fig. 1A**, **STable 1**). These bulk PFC markers include genes identified by more recent studies such as *CBLN2*^17^, as well as genes that represent a variety of functions, including cell cycle regulation, scaffolding and membrane proteins, and regulators of neurogenesis (**STable 1**). A parallel analysis was also performed to isolate markers of the developing visual cortex (V1) which is molecularly distinct to the PFC during development^14^ identifying well-established V1 determinants such as *NR2F1*.

**Fig. 1.**
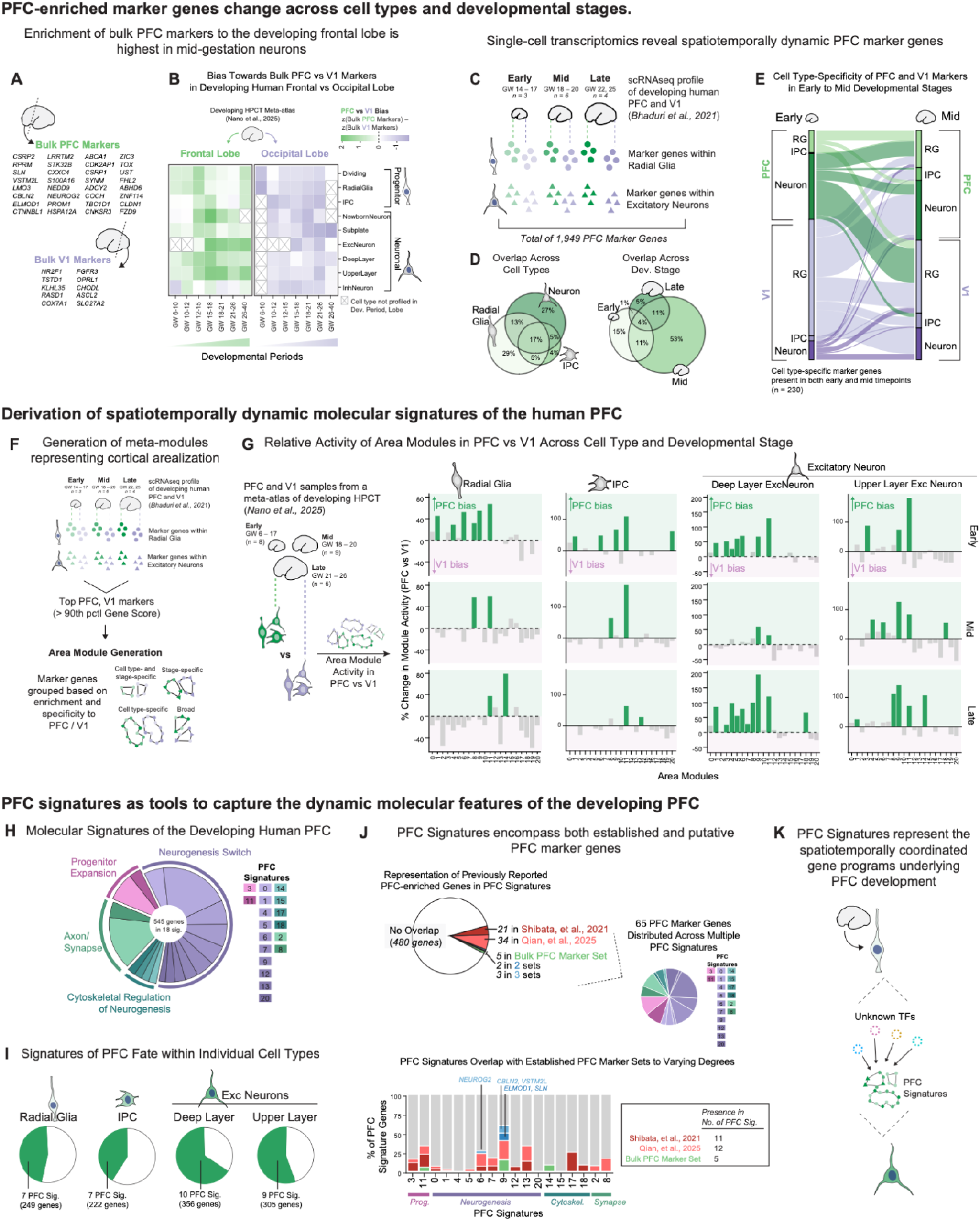
Molecular signatures of the developing prefrontal cortex. **A)** Marker genes for the developing human prefrontal cortex (PFC) vs visual cortex (V1) were identified from a previously published, bulk transcriptomic profile of second-trimester primary human cortex (Miller et al., 2014) and cross-validated in analogous single-cell datasets (Bhaduri et al., 2021) to generate “bulk” PFC and bulk V1 marker sets. **B)** Activity of bulk PFC and V1 marker sets were scored in frontal and occipital lobe samples in a single-cell meta-atlas of developing human primary cortical tissues (HPCT; Nano et al., 2025). Heatmap displays the bias toward bulk PFC versus V1 marker activity across cell types and developmental periods in frontal and occipital lobe dissections. Color intensity reflects the difference between z-scored bulk PFC marker activity and z-scored bulk V1 marker activity, with green indicating greater PFC marker activity. Crossed cells indicate cell types not profiled in the indicated developmental period or lobe. The frontal lobe displays a slight bias toward PFC marker activity, but enrichment of bulk PFC markers in the frontal lobe is dynamic across development and is not a consistent signifier of PFC identity across all stages. **C)** Single-cell transcriptomic profiles of the PFC reveal spatiotemporally dynamic marker genes. Marker genes were identified from a single-cell dataset of the developing human PFC and V1 (Bhaduri et al., 2021) spanning early (GW 14–17, n = 3), mid (GW 18–20, n = 6), and late (GW 22, 25, n = 4) developmental stages. Marker genes were identified within radial glia and excitatory neurons separately, yielding a total of 1,949 PFC marker genes. **D)** Venn diagrams showing the overlap of PFC marker genes across cell types (left) and developmental stages (right), illustrating that the majority of PFC marker genes are specific to a single cell type or developmental stage. **E)** Alluvial plot showing the cell type-specificity of PFC and V1 marker genes in early and mid developmental stages. Cell type-specific marker genes present in both early and mid timepoints (n = 230) are shown, illustrating the conservation and divergence of areal marker gene identity across cell types and developmental stages. **F)** Meta-module analysis to identify dynamic molecular signatures of PFC identity. The meta-module approach (Nano et al., 2025) was applied to the single-cell markers described in Fig. 1C, filtered to the top 90th percentile by gene score, a metric of enrichment and specificity to the PFC vs V1. Meta-module generation grouped these marker genes into modules based on their PFC/V1 enrichment across cell types and developmental stages, yielding 21 modules. **G)** Activity of the 21 area modules generated in Fig. 1F in PFC versus V1 samples from the developing HPCT meta-atlas (Nano et al., 2025) across early (GW 6–17, n = 8), mid (GW 18–20, n = 9), and late (GW 21–26, n = 6) developmental stages. Bar charts show, for each cell type (column) in each developmental stage (row), the relative activity of all 21 area modules in PFC versus V1. Only statistically significant changes are shown (p < 0.05, two-sided Wilcoxon test). Data presented as percent increase or decrease in module activity, relative to that of V1. Green bars indicate PFC bias; purple bars indicate V1 bias. The bias of any given module to the PFC vs the V1 changed across cell types and developmental stages, reflective of the dynamic nature of PFC molecular identity.

To assess these bulk PFC and bulk V1 markers in a broader array of datasets, we leveraged our recently published meta-atlas of single-cell profiles of the developing human cortex^24^ and calculated the difference in bulk marker set activity in samples from the frontal vs occipital lobe. As expected, in the frontal lobe samples, the bulk PFC marker genes were slightly more active than the V1 markers; however this enrichment for bulk PFC markers is dynamic (**Fig. 1B**). Bulk PFC markers are therefore not a consistent signifier of PFC identity, but rather a group of genes that are enriched in a temporal- (primarily GW 12–18) and cell type-specific (primarily neurons) manner. These results are consistent with the fact that our bulk PFC marker set originated from datasets spanning the second trimester^16^. Our analyses therefore indicate that while previously described PFC marker genes can represent a subset of PFC molecular identity, additional gene signatures may exist that more fully capture the heterogeneity and dynamics of the emerging PFC.

To resolve these additional signatures, we leveraged a single-cell transcriptomic dataset of the human PFC and V1 across peak stages of neurogenesis^14^. The resolution of this dataset has enabled the identification of 1,949 PFC markers: genes that are enriched in the PFC compared to the V1, but only within specific cell types at certain developmental stages (**Fig. 1C**). Only a small proportion of these marker genes (4-17%) are consistently enriched in the PFC throughout neuronal differentiation and cortical development (**Fig. 1D**), with several genes displaying large shifts in their cell type- and area-specificity (**Fig. 1E**).

We consolidated these single-cell PFC marker genes into coherent molecular signatures with gene co-expression network analysis. These approaches have enabled our previous work^24^ as well as others^27,28^ to observe changing expression patterns and cell-type enrichments across developmental time and maturation trajectories. Previously, we have published an analytical approach that generates co-expression networks from multiple individuals, designed to overcome technical variation inherent in human samples^24^. This “meta-module” approach is therefore well poised to unify signatures across single-cell samples of human cortical areas at multiple developmental timepoints.

Using our meta-module approach, we consolidated previously identified single-cell PFC markers based on their enrichment to the PFC across cell types and throughout development, yielding 21 modules (**Fig. 1F**, **STable 2**). We then narrowed these modules further, scoring each for their relative activity in PFC versus V1 across the cell types and developmental timepoints of our single-cell, meta-atlas of the developing human cortex^24^ (**Fig. 1G, STable 3**). Any module significantly more active in the PFC across any combination of cell type and developmental stage was considered a signature of molecular PFC fate, resulting in 18 PFC signatures with 545 genes represented (**Fig. 1H-I**). Gene Ontology (GO) term enrichment analysis and literature review showed that broadly these signatures represent developmental processes related to expansion of the progenitor pool, neurogenesis, and synaptic and cytoskeletal aspects of neuronal identity (**SFig. 1, STable 4-5**).

Compared to previous datasets, 65 of the genes in our PFC signatures were previously reported in both the bulk PFC marker set and other previously reported marker genes from single-cell analyses (**Fig. 1J**). Established PFC marker genes were present across 15 of the 18 PFC signatures, with the majority of these established PFC marker genes distributed across multiple PFC signatures. Of note, one neurogenesis-related PFC signature (signature 9) showed the greatest enrichment of previously reported PFC markers (**Fig. 1J**). Given the enrichment of PFC signature 9 in mid-gestation IPC and neurons (Fig. 1G), it is likely that this signature is among the most upregulated at the timepoints of human cortical development most physically accessible for transcriptomic profiling. Our newly defined PFC signatures thus show a broader range of PFC-enriched genes than previous studies, likely because they become enriched in specific cell types and at particular developmental stages. Thus, the PFC signatures here encompass previously identified molecular markers of PFC identity and a broader universe of PFC gene expression in human cortical development (**Fig. 1K**).

### Functional interrogation of intrinsic PFC determinants in HPCT ex vivo cultures

We next examined whether the precise, spatiotemporal coordination of these PFC signatures can be governed intrinsically, independently of extrinsic stimuli. To identify the TFs that are sufficient to activate PFC signatures, we leveraged the recent success of pooled CRISPR activation screening approaches. Pooled CRISPR screening has emerged as a powerful, high-throughput platform to interrogate the molecular players of complex cell fates, with recent examples in cerebral organoid models examining earlier timepoints of telencephalic regionalization^20,29^. More recent screens have deconstructed radial glia lineage trajectories in HPCT *ex vivo* cultures^30^, demonstrating the feasibility of conducting these screens in a highly physiologically relevant model of the developing human cortex^31^.

We therefore conducted our screens in HPCT *ex vivo* cultures, curating screen targets from the spatiotemporally dynamic PFC markers reported in single-cell datasets^14^ (**Fig. 2A**). To maximize the inclusion of genes sufficient to change cell fates, our curation was not limited to our 545 PFC signature genes and instead drew from the complete set of 1,949 single-cell PFC marker genes (**Fig. 2A**). From these PFC marker genes, we selected TFs whose expression levels in hCOs were comparable to or lower than those in primary tissue^14,21^, preserving the opportunity for subsequent mechanistic validation in hCOs. Our final set of 35 screening targets include both well-established regulators of neurogenesis and neuronal differentiation and 14 genes with limited known association with cortical development (**Fig. 2B, STable 6**).

**Fig. 2.**
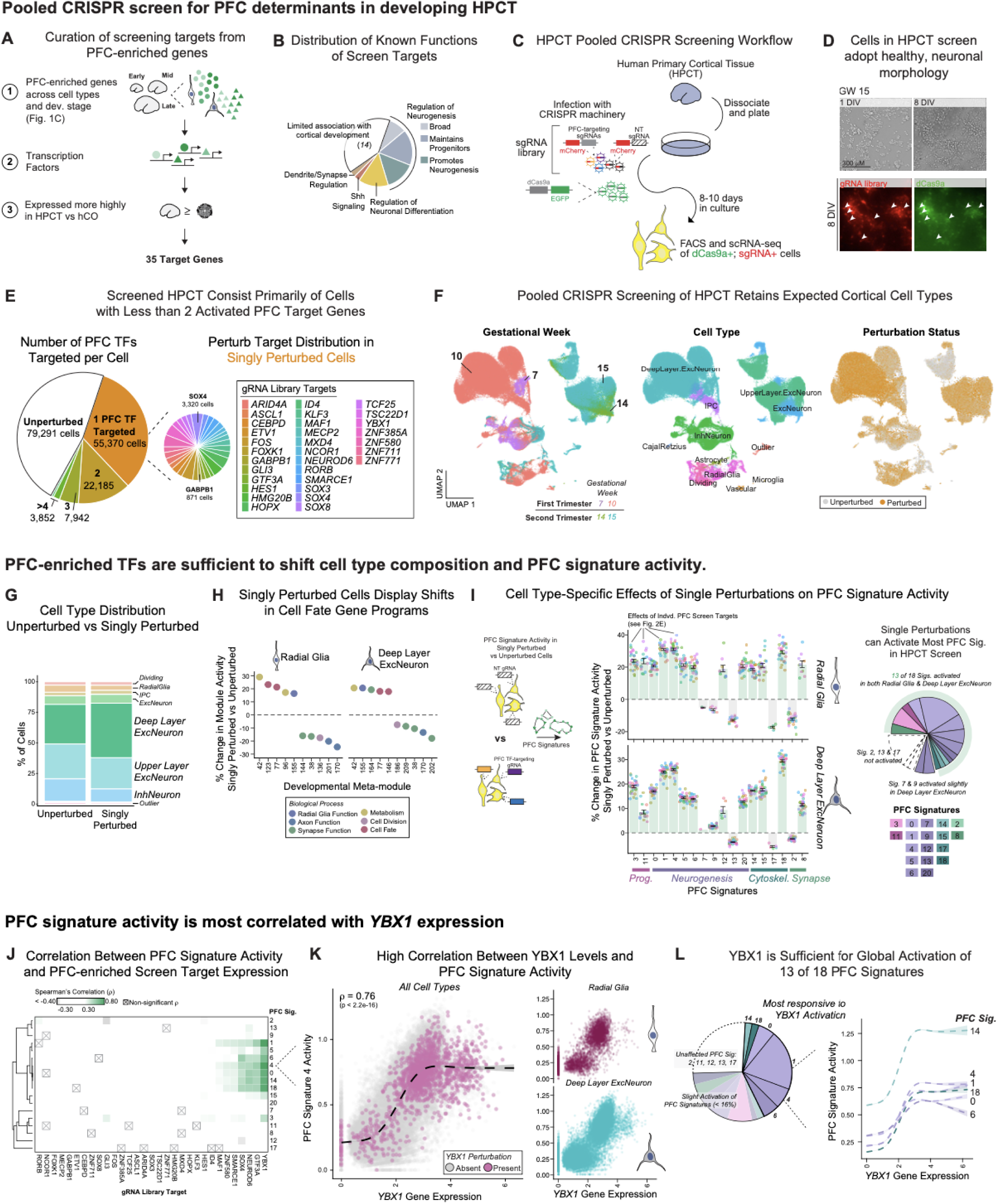
Pooled CRISPR activation screening in human primary cortical tissue (HPCT) identifies transcriptional regulators sufficient to activate PFC signatures. **A)** Screen targets were selected from among the PFC and V1 marker genes used as input for PFC signature generation (Bhaduri et al., 2021). Genes with known roles in transcriptional regulation (e.g. inclusion in ChEA Transcription Factor Dataset) were isolated from these PFC-enriched genes, and, of these, genes expressed at greater levels in HPCT versus human cortical organoids (hCOs) were prioritized. These criteria resulted in 35 PFC screen target genes. **B)** Pie chart displays the known functions among the 35 PFC-enriched transcriptional regulators selected for pooled CRISPR activation screening. Collectively, these targets represent genes with well-established roles in maintaining progenitors, promoting neurogenesis, dendrite/synapse regulation, and regulation of neuronal differentiation, and 14 genes with limited known association with cortical development. **C)** HPCT pooled CRISPR activation screening workflow. HPCT cultures derived from four individuals spanning first trimester (GW 7 and GW 10) and second trimester (GW 14 and GW 15) were dissociated, plated as monolayers, and infected with lentiviral CRISPR activation components 24 hours after plating. These components included a catalytically dead Cas9 fused to a VP64 activator domain with an EGFP reporter, and a gRNA library with 7 gRNAs per target gene and 21 non-targeting gRNAs as controls. Following 10 days of culture, mCherry+/EGFP+ cells were isolated by FACS and captured for scRNA-seq. **D)** Representative live-cell brightfield and fluorescence images of HPCT cells throughout the screening workflow. Top, HPCT cells display healthy cell morphology throughout the 10-day culture period, with cells proliferating and extending processes. Bottom, cells express mCherry and EGFP fluorescent reporters, indicative of infection with the PFC screen gRNA library and dCas9a, respectively. White arrows, mCherry+; GFP+ cells. Scale bar, 300 μm. **E)** Left pie chart displays the composition of our perturbational dataset, consisting of 79,291 unperturbed cells and 55,370 “singly” perturbed cells that harbor the activation of only one PFC screen target. Singly perturbed cells were distributed across 31 PFC target genes that were effectively evaluated in these screens (right pie chart). An additional 33,979 cells harbored multiple perturbations, consisting mostly of doubly perturbed cells. **F)** UMAP of the HPCT perturbational dataset, with cells colored by gestational week (left), annotated cell type (middle), and presence of effective targeting gRNA (perturbation status, right). Radial glia, deep layer neurons, and other cortical cell types were identified, with progenitors intermixing across all samples and deep vs upper layer neuronal subtypes segregating based on gestational week. Expected cortical cell types were present across all perturbation conditions. **G)** Stacked bar plot of cell type composition in unperturbed and singly perturbed cells, showing a shift toward deep layer neuronal cell fates as a result of PFC screen target activation. **H)** Individual PFC screen targets were sufficient to activate previously reported gene programs related to cortical cell fate, neurogenesis, and metabolism (Nano et al., 2025). Dot plots show, within radial glia and deep layer neurons, the percent change in module activity relative to unperturbed cells. Only statistically significant effects are shown (p < 0.05, two-sided Wilcoxon test). **I)** For each PFC screen target, statistically significant effects on PFC signature activity relative to unperturbed were calculated (p < 0.05, two-sided Wilcoxon test). Analysis was conducted specifically within radial glia (top) or deep layer neurons (bottom). Percent-change in signature activity induced by each screen target shown as individual dots, and bar and error bars represent the average percent-change across all screen targets on the indicated signature. Only significant changes in PFC signature activity are displayed (p < 0.05, two-sided Wilcoxon test). The effects of these perturbations differ slightly between radial glia and deep layer neurons, demonstrating the dynamic nature of PFC molecular fate. These results are summarized in the pie chart (right), showing that across both cell types, 13 of the 18 PFC signatures could be activated by at least one putative PFC transcriptional regulator. **J)** Correlation between PFC signature activity and expression levels of PFC screening targets, calculated within cells that were either unperturbed or singly perturbed for the indicated screening target (columns). Eleven PFC screen targets displayed strong positive correlation (ρ > 0.3) with at least one PFC signature. Only statistically significant measurements shown (p < 0.05, Spearman’s rank correlation test). **K)** YBX1 emerges as the PFC screen target most strongly correlated with PFC signature activity (ρ = 0.76, p < 2.2e-16). Left dot plot shows YBX1 gene expression and activity of PFC signature 4 in all cell types that are either unperturbed (grey dots) or harbor exclusively a YBX1 perturbation (pink dots). Dashed line represents estimated smoothed conditional means (general additive model), with 95% confidence interval. Right dot plots show the analogous data for radial glia and deep layer excitatory neurons. **L)** YBX1 is sufficient for the global activation of 13 of 18 PFC signatures. Percent-change on PFC signatures was calculated between exclusively YBX1-perturbed cells versus unperturbed cells. There was high correlation between YBX1 levels and 6 of these PFC signatures (signatures 0, 1, 4, 6, 14, 18), representing the signatures most sensitive to YBX1 activation. PFC signatures 2, 11, 12, 13, and 17 were unaffected.

We screened these putative PFC determinants in *ex vivo* HPCT cultures derived from four individuals (**Fig. 2C**): two representing first trimester (GW7 and 10) and another two from second trimester stages during peak neurogenesis (GW14 and 15). For each sample, cortical tissue was dissociated and plated as a monolayer as previously described^31^, and infected with lentiviral CRISPR activation (CRISPRa) components 24 hours after plating. These components included a catalytically dead Cas9 tagged to an activator domain with an EGFP reporter and a gRNA library with 7 gRNAs for each of the 35 target genes and 21 non-targeted gRNAs as controls (see Methods). HPCT cells maintained healthy morphology throughout 8–10 days of culture, proliferating and extending processes as expected (**Fig. 2D, SFig. 2A**). Following this period, cells expressing CRISPRa components were isolated by fluorescence-activated cell sorting (FACS) and captured for single-cell transcriptomics (scRNA-seq).

We confirmed the cortical identity of these HPCT samples by projecting early-stage samples onto a first-trimester brain region reference^32^ and mid-stage samples onto a second-trimester cortical reference^14^ (**SFig. 2B**). To determine which PFC screen targets were effectively activated in individual cells, we developed a rigorous computational pipeline for single-cell gRNA detection. Recognizing that gRNA efficacy is highly context-dependent^33^, we sought to optimize the detection of perturbed cells by employing the most stringent negative controls available among the non-targeting gRNAs. For each of the 35 PFC screen target genes, our pipeline identified: (1) the subset of non-targeting gRNAs with no detectable off-target effects on that gene’s expression; (2) the targeting gRNAs that significantly upregulated target gene expression relative to this gene-specific negative control population; and (3) the cells harboring these valid targeting gRNAs at sufficient abundance to be classified as perturbed (**SFig. 2C-E**, **STables 7-9**, see Methods). These analyses enabled the classification of each cell as unperturbed, singly perturbed, or multiply perturbed, providing a high-confidence perturbational dataset for downstream analyses.

These screens produced a perturbational dataset consisting of a negative-control population of 79,291 “unperturbed” cells, and 55,270 “singly perturbed” cells harboring the activation of only one PFC screen target, with cells evenly distributed across the 31 PFC target genes that were effectively screened in these experiments (**Fig. 2E**). An additional 33,979 cells harbored multiple perturbations, consisting mostly of “double-perturbed” cells.

To evaluate how these perturbations change cell fate, we first confirmed the presence of expected cortical cell types in this perturbational dataset (**Fig. 2F, SFig. 3A, STable 10-11**). Our HPCT screens also retained expected cell type shifts across development: while we observed higher transcriptomic similarity within progenitor cells and inhibitory neurons across all samples, excitatory neurons harboring deep layer identities were predominant in first-trimester samples and more mature, upper layer identities were more prevalent the second trimester. In both developmental stages, perturbed cells were present in all neural cell types (**SFig. 3B**). We found that the activation of PFC screening targets resulted in a shift towards deep layer neuronal cell fates: specifically, deep layer neurons are expanded in the singly perturbed population vs unperturbed cells (**Fig. 2G, SFig. 3C**). This was expected because many of our screening targets are known to regulate neurogenesis, and indeed, singly perturbed cells displayed significant activation of cell fate, neurogenesis, and metabolism gene programs relative to unperturbed controls (**Fig. 2H, STable 12**).

### Individual PFC TFs are sufficient to activate PFC signatures

Given that individual PFC TFs were sufficient to alter cell fates in these HPCT cultures, we next evaluated the effects of these screen targets on our 18 PFC signatures (**Fig. 2I, SFig. 3D, STable 13**). We focused particularly on the effects of PFC TFs within cell types, e.g. radial glia and deep layer neurons: most PFC signatures were activated by at least one PFC TF (**Fig. 2I**), with PFC TFs displaying concordant effects across these signatures. Signatures that were resistant to intrinsic activation reflect potential aspects of extrinsic communication lost in our *ex vivo* cultures. For example, our PFC screen targets failed to activate PFC Signature 17, which is associated with radial glia delamination and cytoskeletal organization (see **STable 5**). Six PFC screen targets were themselves members of a PFC signature, and, of these, all but one were sufficient to activate their respective “home” signature (**SFig. 3E**). The exception was NEUROD6, a neurogenesis driver whose home signature (PFC Signature 13) is associated with axon formation (**STable 5**) and was not activated by any of our PFC screen targets. Though the synergistic effects of these TFs was not the focus of this screen, we also observed that the activation of PFC TF pairs also showed significantly additive activation of PFC signatures (**SFig. 3F-H**). Together, these analyses demonstrate that diverse PFC-enriched TFs are individually sufficient to activate discrete subsets of the PFC gene expression universe.

### YBX1 emerges as a key dose-dependent determinant of the PFC

We hypothesized that the necessary and sufficient levers of the dynamic domino effect observed in the developing human cortex^14^, would confer sensitive, dose-dependent responses on PFC signature activity. We looked for such factors among our PFC screening targets, evaluating the global correlation between PFC signature activity and expression levels of PFC TFs **(Fig. 2J, STable 14)**. Seven PFC screen targets were strongly correlated with at least one PFC signature, with the key standout being the neurogenesis regulator *YBX1* (**Fig. 2K**).

YBX1 has well-established roles in maintaining radial glia progenitor pools in the developing mouse cortex^26^ and promoting forebrain specification^25^, with disease relevance as an oncogene in several brain tumors^34,35^, including glioblastoma^36^. Recent mechanistic studies allude to the role of YBX1 in binding to and modulating the activity of chromatin regulators^25^, stabilizing of synaptic mRNA transcripts^37^, and regulating RNA splicing via physical interaction with MECP2, itself a target in our screens^38^. In this activation screen, *YBX1* is sufficient for the global activation of 13 of 18 PFC signatures, with a subset of 6 signatures being most responsive to *YBX1* activation (**Fig. 2L**). These findings point to the ability of YBX1 to intrinsically modulate PFC fate, most directly on 6 of the 18 PFC signatures. By providing functional evaluation of YBX1 and 30 additional PFC TFs, our pooled CRISPR screens in HPCT *ex vivo* cultures establish a perturbational reference dataset of PFC determinants and demonstrate the contribution of intrinsic cues on PFC fate specification.

### Benchmarking the activity of PFC determinants in hCO

While our HPCT pooled CRISPR screens establish a valuable perturbational dataset, hCOs offer a more accessible platform for characterizing the mechanisms of action of these PFC determinants. Recent work has demonstrated the versatility of hCOs to screen for regulators of several stages of neurodevelopment, including early regionalization^29^ and fate specification^20^. However, the study of areal patterning in hCOs is challenged by the reports of the stochasticity and lack of spatial area organization in these systems^21^. Rigorous benchmarking of how PFC TFs operate in hCOs relative to HPCT is therefore a prerequisite for deploying these models as mechanistic testing grounds for intrinsic PFC determinants.

We therefore screened our 35 PFC screen targets in hCOs. To maximize the efficiency of perturbations, we applied the reaggregation and infection approach that we have previously implemented, inspired by chimeroid systems^24,39^ (**Fig. 3A**). CRISPRa machinery was delivered to cortical organoids at about 3 weeks of culture, and perturbed cells were isolated by FACS and analyzed with scRNA-seq (**SFig. 4A-B, STable 15-16**). Analysis of the cells passing rigorous QC showed that the majority of cells in these screened organoids retained cortical identity (**Fig. 3B, SFig. 4C, STable 17**), though we observed that dCas9 expression resulted in an increased rate of non-cortical cells that was potentiated by 6 individual PFC TFs (**SFig. 4D**). Notably, activation of PFC screen targets broadly decreased the proportion of non-cortical cell types and increased deep layer neuronal subtypes relative to unperturbed cells (**Fig. 3C**), mirroring observations in our HPCT screens. For subsequent analyses, we proceeded to focus on the cells annotated as cortical.

**Fig. 3.**
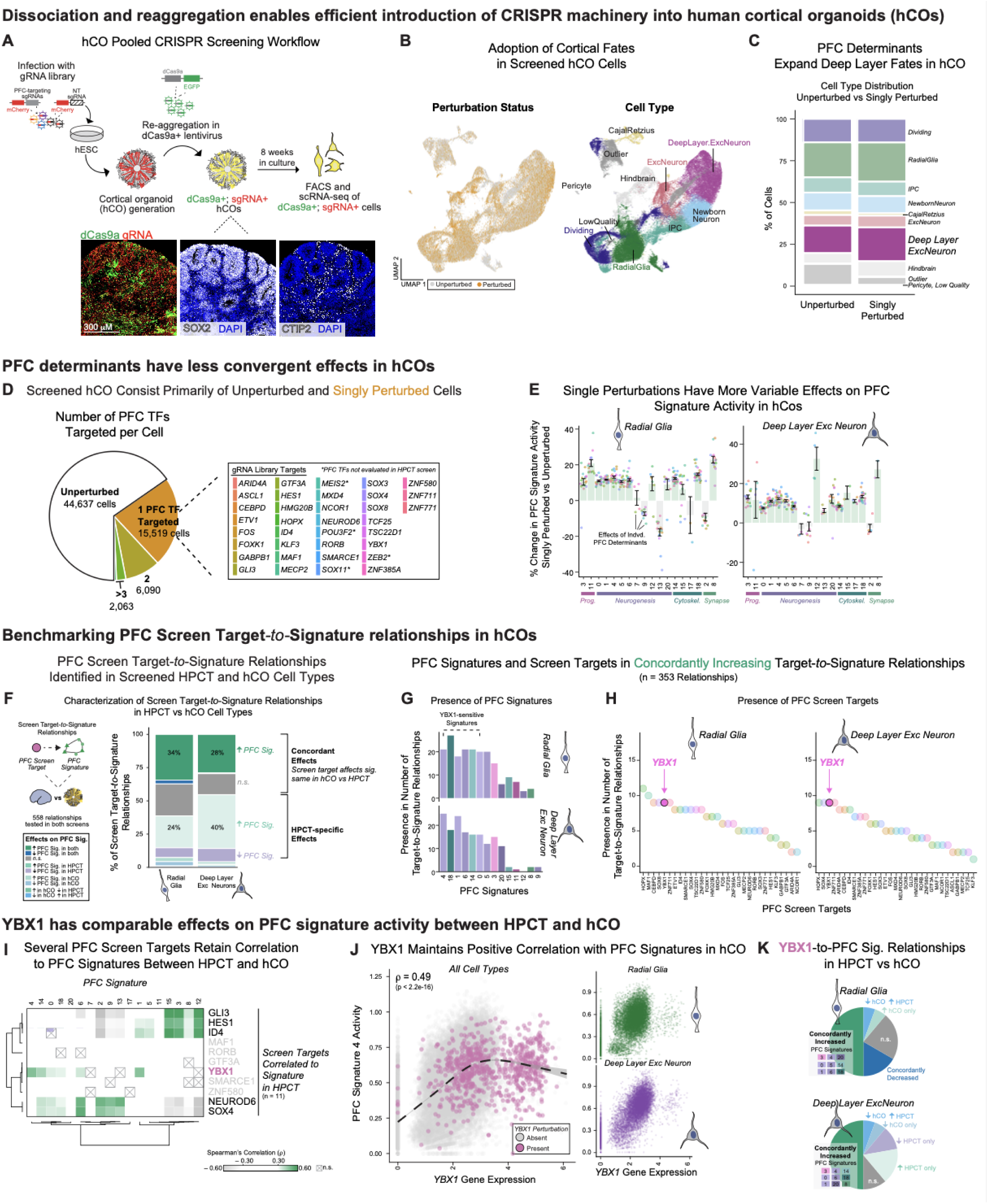
Benchmarking human cortical organoids (hCOs) as a pooled CRISPR screening platform. **A)** Schematic of the hCO pooled CRISPR activation screening workflow. Human embryonic stem cells (hESCs) were infected with the same mCherry+/gRNA library of PFC screen targets as used in HPCT screens (Fig. 2) and used to generate cortical organoids. After 18 days of culture, organoids were dissociated and re-aggregated in the presence of EGFP+/dCas9a+ lentivirus (Anton-Bolanos, et al., 2024; Nano et al., 2025). Organoids were grown for an additional 8 weeks, at which time mCherry+/EGFP+ cells were isolated by FACS and captured for scRNA-seq. Representative immunofluorescence images show EGFP (dCas9a) and mCherry (gRNA) expression, as well as SOX2 (progenitors), and CTIP2 (deep layer neurons) in screened hCOs, corroborating infection of the CRISPR machinery and retention of cortical identity. Scale bar = 300 μm. **B)** UMAP of screened hCO cells colored by perturbation status (left) and annotated cell type (right), demonstrating that the majority of cells in screened organoids retained cortical identity, including expected progenitor, neuronal, and non-neuronal cell types. **C)** Stacked bar plot of cell type distribution in unperturbed versus singly perturbed cells in screened hCOs. Mirroring HPCT screens, activation of PFC determinants results in a slight expansion of Deep Layer Excitatory Neurons in the singly perturbed population. **D)** Screened hCOs consist primarily of unperturbed (44,637 cells) and singly perturbed cells (15,519 cells harboring activation of 1 PFC TF), with 6,090 doubly perturbed cells and 2,063 cells harboring more than 3 perturbations. The full list of PFC targets effectively screened in hCOs are listed, including four library targets that were not effectively evaluated in the HPCT screen (*MEIS2*, *POU3F2*, *SOX11*, *ZEB2*, marked with an asterisk). **E)** Single perturbations have more variable effects on PFC signature activity in hCOs compared to HPCT. Bar and error bars represent the average percent change in PFC signature activity in singly perturbed versus unperturbed cells, displayed separately for radial glia and deep layer excitatory neurons across all 18 PFC signatures (grouped by biological category). Dots show the effects of individual PFC determinants. Only statistically significant effects are shown (p < 0.05, two-sided Wilcoxon test). **F)** Characterization of PFC Screen Target-to-Signature relationships between HPCT and hCO cell types (558 relationships tested in both screens). Stacked bar chart displays the proportion of relationships in each category: e.g. concordantly increased in both systems; concordantly decreased; HPCT-specific and hCO-specific effects. In radial glia, 34% of relationships resulted in PFC signature activation in both HPCT and hCO, while 24% only resulted in signature activation in HPCT. In deep layer excitatory neurons, 28% were concordantly increased and 40% demonstrated HPCT-specific activation. These comparisons suggest that extrinsic cues present in primary tissue but absent from hCOs may be required for the convergent effects of PFC determinants observed in HPCT. **G)** Across both cell types, 353 PFC Screen Target-to-Signature relationships resulted in signature activation in both HPCT and hCO. Bar charts show the number of concordantly increasing relationships that involve each PFC signature (x-axis), stratified by radial glia and deep layer excitatory neurons. YBX1-sensitive PFC signatures are most frequently represented in concordantly increasing relationships across both cell types. **H)** YBX1 is among the screen targets with most frequently involved in concordantly increasing relationships. For both radial glia (left) and deep layer neurons (right), dot plots display the number of concordantly increasing Target-to-Signature relationships that involve each PFC screen target. **I)** Correlation between PFC signature activity and expression levels of PFC screening targets: specifically, the 11 screen targets correlated to at least one signature in HPCT screens (Fig. 2K). Of these, 7 targets retain significant correlation to PFC signatures in hCO. Only statistically significant measurements shown (p < 0.05, Spearman’s rank correlation test). **J)** YBX1 maintains a positive correlation with PFC signature activity in hCOs (ρ = 0.49, p < 2.2e-16, Spearman’s rank correlation test). Scatter plots display PFC Signature 4 activity versus YBX1 gene expression across all cell types (left), and separately within radial glia and deep layer excitatory neurons (right), with YBX1 perturbation status indicated (pink dots; grey, unperturbed). Dashed line represents estimated smoothed conditional means (general additive model), with 95% confidence interval **K)** Pie charts comparing YBX1-to-PFC Signature relationships between HPCT and hCO, separately for radial glia and deep layer excitatory neurons. Data shown are the proportion of relationships that are concordantly increased, concordantly decreased, or display effects specific to hCO vs HPCT screens. Half of the YBX1-to-PFC Signature relationships result in signature activation in both systems, highlighting the robustness of YBX1’s effects across experimental models.

Pooled CRISPR screening approach in the hCO system once again resulted in a majority of unperturbed and singly perturbed cells (**Fig. 3D, SFig. 4E-F, STable 18**). However, unlike in our HPCT screens, the effects of individual PFC TFs on PFC signatures were more variable and less convergent in terms of both magnitude and directionality (**Fig. 3E, SFig. 4G, STable 19**). The hCO system may therefore lack regulatory mechanisms present in *ex vivo* HPCT cultures that restrict the activity of PFC determinants into the more convergent effects we observed in HPCT.

### Key PFC Gene Signatures and YBX1 dose dependency are retained in hCO

These initial analyses suggest that PFC screen targets vary in their ability to be modeled in the hCOs. To isolate the PFC screen targets that can be accurately studied in hCOs, we systematically evaluated “PFC Screen Target-*to*-Signature” relationships: how the activation of each screen target gene regulates each PFC signature (**Fig. 3F**). We accounted for cell type-specific effects by conducting these hCO vs HPCT comparisons within radial glia and deep layer neurons. While as much as 40% of these relationships resulted in activation exclusively in HPCTs, we observed that at least 25% of PFC Screen Target-*to*-Signature relationships resulted in PFC signature activation in both systems.

We hypothesized that these concordantly increased relationships represent the PFC determinants and signatures that can be most robustly examined in hCO and HPCT. Interestingly, across both radial glia and deep layer neurons, these high-value PFC Screen Target-*to*-Signature relationships most frequently involve the same set of 6 YBX1-sensitive PFC signatures first identified in our HPCT screen (**Fig. 3G**). When we performed the inverse analysis, we found that YBX1 is among the most frequent screen targets involved in these concordantly increasing relationships: specifically, in deep layer neurons, YBX1 is among the screen targets that can activate the most PFC signatures in both hCO and HPCT (**Fig. 3H**). In addition, YBX1 still retained its dose-dependent effects on these 6 YBX1-responsive signatures in the hCO (**Fig. 3I-J, SFig. 4H-I, STable 20**). We therefore focused again on YBX1, comparing its effects on cell fate across both screening platforms (**STable 18**). We observed a substantial overlap in the developmental gene programs changed by YBX1 in hCOs vs HPCT within radial glia (**SFig. 4J**). In deep layer neurons, YBX1 activation in hCOs modulated a subset of the developmental gene programs that it changed in HPCT, with gene modules relating to mature axon and synaptic functions relatively unaffected in the hCO (**SFig. 4K**). However, in both radial glia and deep layer neurons, YBX1 retained the ability to increase 9 PFC signatures in both hCO and HPCT systems (**Fig. 3K**). Together, these analyses identify the subset of intrinsic PFC determinants whose regulatory relationships with PFC signature are preserved in hCOs, including YBX1 and its dose-dependent relationship with 6 PFC signatures.

### YBX1 plays essential roles in PFC signature activation

Our findings that YBX1 is sufficient to activate a subset of PFC signatures accompany its established roles in regulating the chromatin landscape of neuronal progenitors^25^. We therefore tested whether YBX1 is essential for the maintenance of PFC fate and how these functions intersect with its role as a chromatin regulator. We again leveraged our targeted genetic perturbation approaches in chimeroid systems, but now with the introduction of shRNAs to knock-down YBX1 and profile the perturbed cells with single-cell multiomic profiling (**Fig. 4A-B, SFig. 5A-C, STable 21-22**). In line with previous studies^26^, YBX1 depletion in hCOs led to an expansion of deep layer neuronal subtypes (**Fig. 4C, SFig. 5C**) and the activation of developmental gene programs related to synapse function, including a module that we have previously linked to deep layer fates^24^ (**Fig. 4D, STable 23**).

**Fig. 4.**
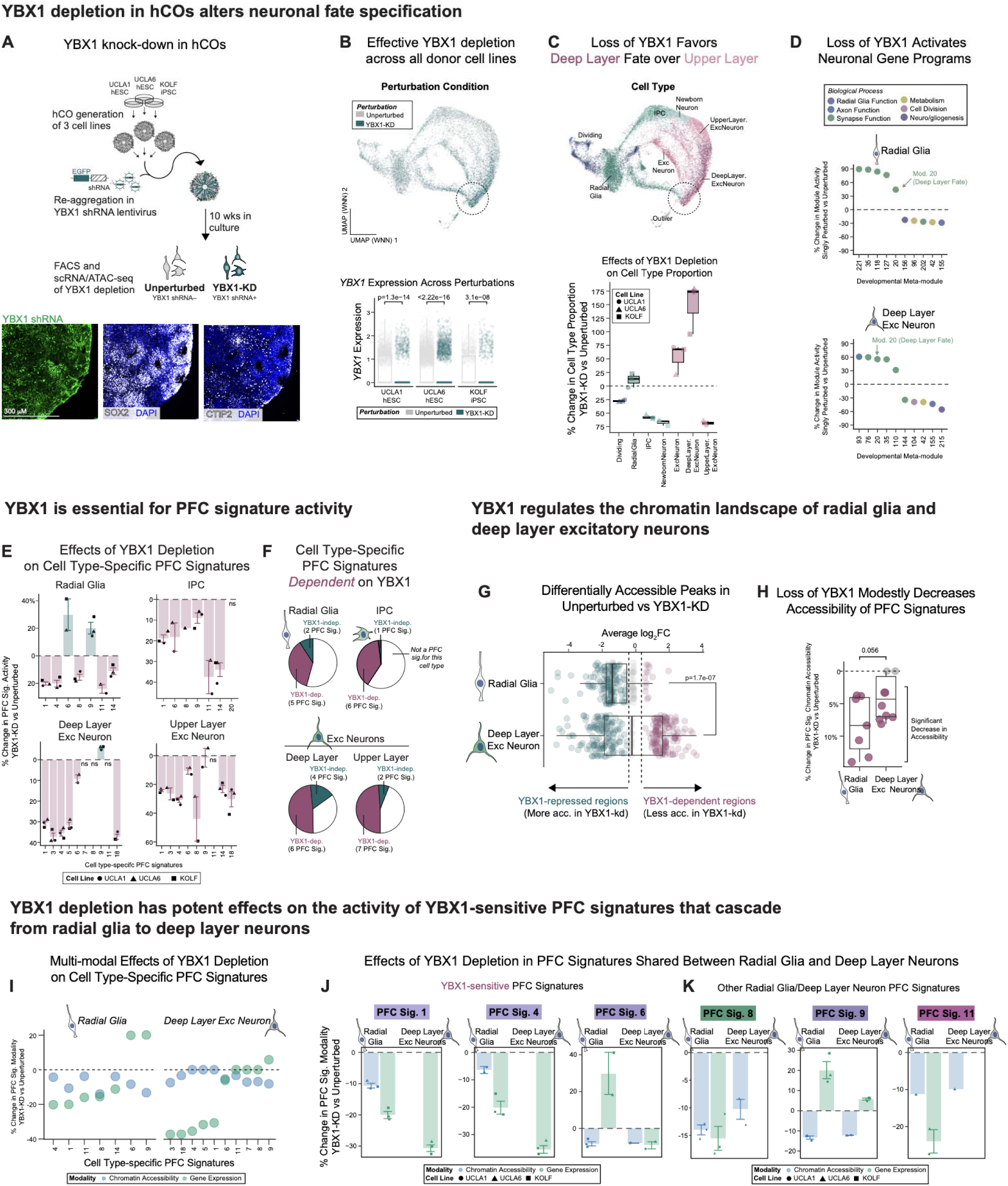
YBX1 plays essential roles in the epigenetic and transcriptional regulation of molecular PFC fate. **A)** Schematic of the YBX1 knock-down (KD) workflow in hCOs. Three stem cell lines (UCLA1 hESC, UCLA6 hESC, KOLF iPSC) were infected with a YBX1-targeting shRNA (EGFP-labeled). Cortical organoids were generated from each cell line, which, after 18 days, were dissociated and re-aggregated together in the presence of YBX1 shRNA lentivirus (Anton-Bolanos, et al., 2024; Nano et al., 2025). The resulting chimeroids were grown for 8 weeks in culture, at which time EGFP+ (YBX1-KD) and EGFP– (unperturbed) cells were isolated by FACS and captured for single-cell multi-omic profiling (simultaneous scRNA-seq and scATAC-seq). Representative immunofluorescence images of 5-week-old chimeroids show YBX1 shRNA (EGFP, green), SOX2 (DAPI), and CTIP2 (DAPI), confirming cortical identity. Scale bar = 300 μm. **B)** YBX1 expression is significantly and consistently depleted in YBX1-KD cells across all three cell lines. (Top) UMAP projection from both RNA and ATAC modalities (UMAP WNN) of YBX1-KD hCO cells colored by perturbation condition. (Bottom) YBX1 expression in unperturbed versus YBX1-KD cells for each cell line, summarized with boxplots (two-sided Wilcoxon test). **C)** Loss of YBX1 favors deep layer fate over upper layer fate. (Top) UMAP (WNN) projection of cells colored by cell type, with the Deep Layer Excitatory Neuron cluster highlighted. (Bottom) Percent change in cell type proportion in YBX1-KD versus unperturbed cells for each cell type. Individual dots represent each cell line, data summarized by boxplot. Dashed line at 0 indicates no change. **D)** Loss of YBX1 activates neuronal gene programs in both radial glia and deep layer neurons. Dot plots show the percent change in activity of developmental meta-modules (Nano et al., 2025) in YBX1-KD versus unperturbed cells in radial glia (top) and deep layer excitatory neurons (bottom), colored by biological process. Module 20, associated with deep layer fate (Nano et al., 2025), is among the most elevated modules in both cell types. Only statistically significant effects are shown (p < 0.05, two-sided Wilcoxon test). **E)** YBX1 depletion attenuates most PFC signatures across the excitatory neuronal lineage. Bar and error bars show the average percent change in PFC signature activity in YBX1-KD versus unperturbed cells across radial glia, IPC, deep layer excitatory neurons, and upper layer excitatory neurons. Values for each individual cell line shown as dots. Only statistically significant effects are shown (p < 0.05, two-sided Wilcoxon test). **F)** Across the glutamatergic lineage, YBX1 is required for the majority of cell type-specific PFC signatures. Pie charts summarizing the number of cell type-specific PFC signatures that are dependent (pink) versus independent (teal) of YBX1 in each cell type. White indicates signatures that are not enriched in the PFC within the indicated cell type. **G)** YBX1 depletion induces broad shifts in chromatin accessibility, with effects significantly amplifying from radial glia to deep layer excitatory neurons (two-sided Wilcoxon test). Dots represent significantly differentially accessible chromatin peaks between unperturbed and YBX1-KD cells within radial glia and deep layer excitatory neurons, plotted based on fold-change in accessibility (p < 0.05, logistic regression test). Data are colored by YBX1-dependent regions (less accessible in YBX1-KD), YBX1-repressed regions (more accessible in YBX1-KD), and relatively unchanged regions (less than 25% change in accessibility, grey dots). Data summarized by boxplot. **H)** Loss of YBX1 modestly decreases the chromatin accessibility of PFC signatures. The average promoter region accessibility in each PFC signature was calculated per cell. Data show the percent change in this “PFC signature chromatin accessibility” in YBX1-KD vs unperturbed cells. Comparisons were conducted within radial glia and deep layer neurons, focusing solely on the PFC signatures relevant to each cell type. Dots indicate the percent-change in average chromatin accessibility for each cell type-specific signature, summarized by boxplots. P-value calculated by two-sided Wilcoxon test. **I)** YBX1 regulates PFC signatures at both the chromatin and transcriptional level in radial glia, but acts predominantly as a transcriptional regulator in deep layer neurons. Dot plot displays the percent change induced by YBX1-KD in each cell type-specific PFC signature, both in terms of average chromatin accessibility (blue) and gene expression (green). Dashed line at 0 indicates no change. **J–K)** In YBX1-sensitive PFC signatures shared between radial glia and deep layer neurons (J), YBX1 is required to open chromatin in radial glia but shifts to a predominantly transcriptional role in deep layer neurons – a cascade not observed in non-YBX1-sensitive signatures (K). Bar and error bars show, for the indicated PFC signatures, the average percent change in chromatin accessibility (blue) and gene expression (green) induced by YBX1-KD. Effects in radial glia and deep layer neurons are shown, with the values from individual cell lines shown as dots.

In addition to these previously described YBX1 knock-down phenotypes, we observed that YBX1 is essential for PFC fate across the excitatory neuronal lineage: depletion of YBX1 attenuated the expression of most PFC signatures across radial glia, IPCs, deep and upper layer neurons (**Fig. 4E-F, STable 24**).

With confirmation of the transcriptomic necessity of YBX1, we examined its effects in the chromatin accessibility of these cell types. Consistent with its known roles regulating chromatin remodelers^25^, we observe that YBX1 depletion induces broad shifts in chromatin accessibility across both radial glia and deep layer neurons (**Fig. 4G, SFig. 5D-E, STable 25**). The effects of YBX1 knock-down amplify between these cell types, with YBX1 depletion in deep layer neurons impacting a greater number of chromatin regions to greater effect. However, when we focused this analysis on regions associated with PFC signature genes, we observed that loss of YBX1 induced mild effects on PFC signature chromatin accessibility (**Fig. 4H, STable 26**).

To systematically evaluate the transcriptomic and epigenomic roles of YBX1 on PFC fate, we compared the necessity of YBX1 on the chromatin accessibility vs gene expression of PFC signatures (**Fig. 4I, STable 27**). Within radial glia, YBX1 regulates PFC signatures at the level of both modalities to modest and slightly variable effect. In deep layer neurons, YBX1 acts as a potent and predominantly transcriptional regulator of these PFC signatures. We examined these cascading effects on PFC fate more closely by tracking the multi-modal effects of YBX1 on individual PFC signatures across radial glia and deep layer neurons (**Fig. 4J-K**). Six signatures are enriched in both radial glia and deep layer neurons in the PFC. These include 3 of the 6 YBX1-sensitive signatures identified in our HPCT screens: in 2 of these, we observed that YBX1 is required to open the chromatin landscape of radial glia, then shifts to a stronger, exclusively transcriptomic regulator of these signatures in neurons (**Fig. 4J**). These patterns contrast with those observed in PFC signatures that are not sensitive to YBX1 activity (**Fig. 4K**). YBX1 knock-down in hCOs therefore provide a complementary validation of the YBX1 activation phenotypes observed in HPCT and hCOs: YBX1 is necessary and sufficient for PFC signature activity, with particularly sensitive effects on PFC signature 1 and 4 through staggered effects on chromatin accessibility and gene expression.

### YBX1 activates a cohort of PFC determinants

Across screened and targeted perturbation experiments in HPCT and hCO systems, our data show that YBX1 is a sufficient and essential activator of PFC signatures. However, many of the 34 additional gene targets curated for our pooled CRISPR screens have similar roles in neurogenesis and cell fate (**STable 6**) and, as observed in our HPCT pooled activation screen, activate several PFC signatures. Moreover, known interactions exist between these 35 PFC screening targets (**SFig. 6A**). We therefore examined how YBX1 acts in concert with other PFC TFs to orchestrate the spatiotemporally dynamic expression of PFC signatures, leveraging our perturbational datasets in HPCT and hCO.

We first examined whether YBX1 is an essential activator of other PFC screen targets, using our YBX1 knock-down dataset in hCO. YBX1 is necessary for the chromatin accessibility and gene expression of several PFC targets, most notably within radial glia and deep layer neurons (**Fig. 5A**). Again, we observed that the effects of YBX1 knockdown on chromatin accessibility of these PFC-enriched transcriptional regulators diminishes from radial glia to deep layer neurons. Mirroring our observed effects of YBX1 on PFC signatures, YBX1 may therefore transition into a transcriptional activator of PFC TFs in deep layer subtypes.

**Fig. 5.**
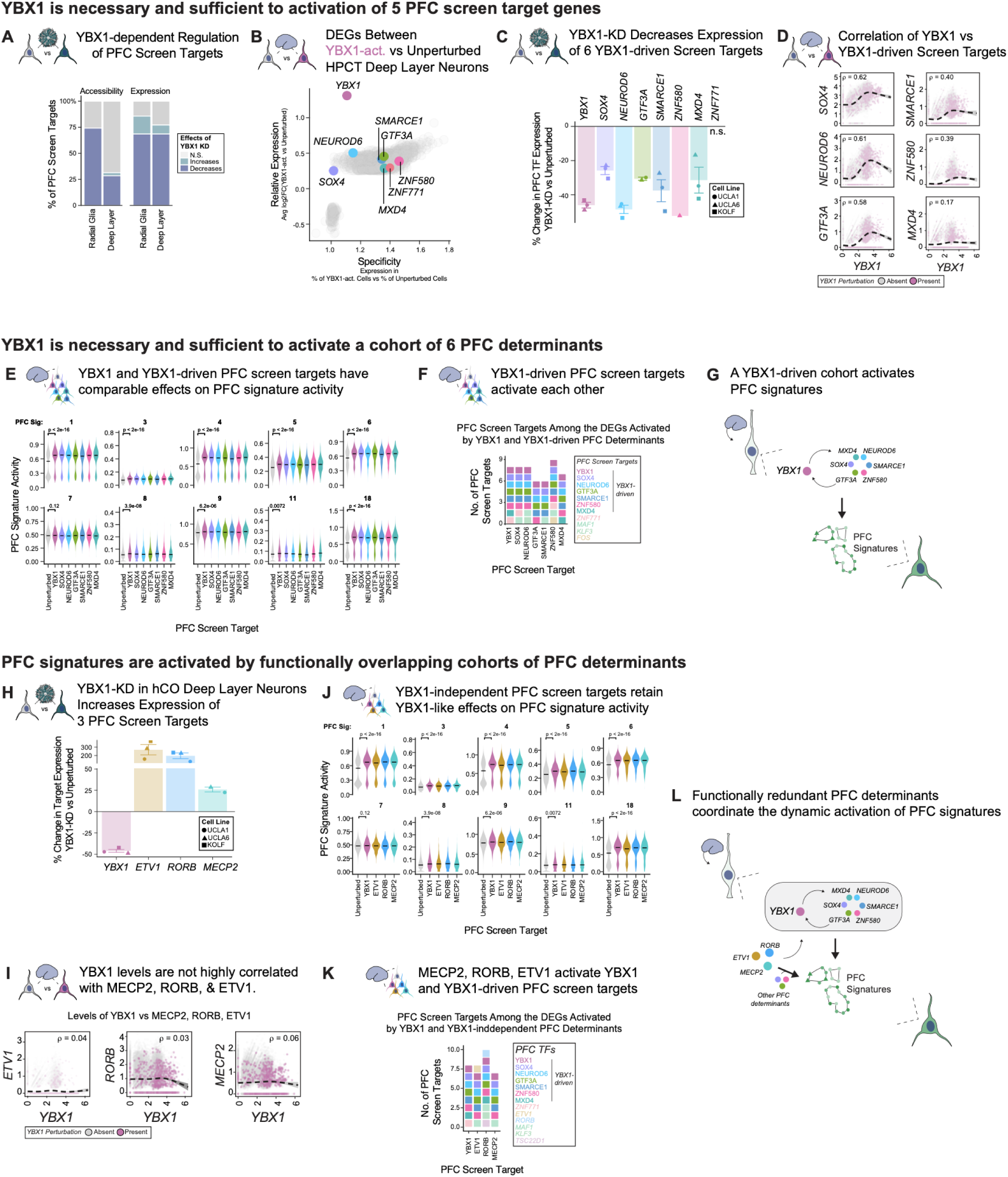
A cohort of YBX1-driven PFC determinants patterns PFC fate in human deep layer neurons. **A)** YBX1 plays an essential role in the regulation of other PFC screen targets. Bar chart shows the proportion of PFC screen targets whose chromatin accessibility and gene expression are significantly increased, decreased, or not significantly changed by YBX1 knock-down. In radial glia, loss of YBX1 decreases both the accessibility of PFC screen target promoter regions and their gene expression levels. In contrast, YBX1 is essential primarily for target gene expression in deep layer neurons. **B)** In the deep layer neurons of our HPCT screens, YBX1 activation is sufficient to upregulate 6 other PFC screen targets. Scatter plot of differentially expressed genes (DEGs) between exclusively YBX1 singly perturbed versus unperturbed HPCT deep layer neurons. DEGs plotted based on their specificity to YBX1-activated populations (x-axis) and fold-change in expression compared to unperturbed cells (y-axis). *YBX1*, *SOX4*, *NEUROD6*, *GTF3A*, *SMARCE1*, *ZNF580*, *ZNF771*, and *MXD4* are among the significantly upregulated genes, identifying a putative YBX1-driven cohort of PFC screen targets. **C)** In our hCO YBX1 knock-down experiments, 6 YBX1-driven screen targets required YBX1 for maximal gene expression. Bar and error bars show the average percent change in gene expression in YBX1-KD versus unperturbed hCO deep layer neurons, with individual cell line values shown as dots. Only statistically significant effects are shown (p < 0.05, two-sided Wilcoxon test). Note that *ZNF711* was not significantly affected by YBX1 knock-down, and was not analyzed further. **D)** Dose-dependent relationship between YBX1 and each YBX1-driven screen target in HPCT deep layer neurons. Scatter plots demonstrate the correlation (ρ) of YBX1 expression versus indicated target genes, with YBX1 perturbation status indicated. Correlations are statistically significant (p < 0.05, Spearman’s rank correlation test). Dashed line represents estimated smoothed conditional means (general additive model), with 95% confidence interval. **E)** YBX1 and the YBX1-driven screen targets activate PFC signatures to comparable levels in HPCT deep layer neurons. Violin plots showing activity of deep layer neuron-specific PFC signatures in unperturbed cells and cells singly perturbed for *YBX1*, *SOX4*, *NEUROD6*, *GTF3A*, *SMARCE1*, *ZNF580*, or *MXD4*. Significance values are indicated above each comparison (two-sided Wilcoxon test). **F)** YBX1-driven screen targets activate each other, consistent with a self-reinforcing regulatory cohort. Stacked bar plot shows the number of PFC screen targets among the DEGs activated by YBX1 and each YBX1-driven PFC determinant in HPCT deep layer neurons. **G)** A model for YBX1-driven PFC fate specification: YBX1 is necessary and sufficient to activate a cohort of PFC-enriched transcriptional regulators (*SOX4*, *NEUROD6*, *GTF3A*, *SMARCE1*, *ZNF580*, *MXD4*) that in turn drive PFC signature activity in deep layer neurons. **H)** In contrast to the YBX1-driven cohort, YBX1 knock-down increases the expression of ETV1, RORB, and MECP2, suggesting these factors operate upstream or in parallel to YBX1. Bar and error bars show the average percent change in PFC screen target expression in YBX1-KD versus unperturbed hCO deep layer neurons for YBX1, ETV1, RORB, and MECP2, with individual cell line values shown as dots. Only statistically significant effects are shown (p < 0.05, two-sided Wilcoxon test). **I)** The absence of meaningful correlation between YBX1 and ETV1, RORB, and MECP2 suggests that these three factors operate independently of YBX1 dosage. Scatter plots showing the correlation of YBX1 expression versus MECP2, RORB, and ETV1 in HPCT deep layer neurons, with YBX1 perturbation status indicated. Dashed line represents estimated smoothed conditional means (general additive model), with 95% confidence interval. **J)** Despite their independence from YBX1 dosage, ETV1, RORB, and MECP2 activate PFC signatures to levels comparable to YBX1 in HPCT deep layer neurons, suggesting functional redundancy. Violin plots showing PFC signature activity in unperturbed cells and cells singly perturbed for YBX1, ETV1, RORB, or MECP2, across deep layer-specific PFC signatures. Significance values are indicated above each comparison (two-sided Wilcoxon test). **K)** ETV1, RORB, and MECP2 activate YBX1 and the YBX1-driven cohort. Stacked bar plot shows the number of PFC screen targets among the differentially expressed genes (DEGs) activated by YBX1 and each of ETV1, RORB, and MECP2 in HPCT deep layer neurons. **L)** Together, our perturbational datasets suggest a framework for coordinated PFC fate specification. ETV1, RORB, and MECP2 act as parallel, functionally redundant PFC determinants that can activate YBX1 and the YBX1-driven cohort, forming a more robust network to coordinate PFC signature activity in deep layer neurons.

To determine which of these downregulated PFC screening targets are most directly regulated by YBX1, we turned to the YBX1-activated deep layer populations from our HPCT pooled activation screen. YBX1 activated only 6 other target genes in our PFC screening library (**Fig. 5B, STable 28**), all of which were also identified as YBX1-dependent (**Fig. 5C**, **SFig. 6B**). Of these, *SOX4*, *NEUROD6*, and *GTF3A* were the most sensitive to YBX1 activity, with expression levels that were tightly correlated with that of *YBX1* (**Fig. 5D**). These comparative analyses therefore show that YBX1 is necessary and sufficient to activate a cohort of 6 PFC TFs.

Our HPCT pooled activation screen dataset enabled us to explore whether these YBX1-driven TFs transduce YBX1-driven PFC fate in deep layer neurons. Five of the six genes in this cohort were sufficient to drive similar effects on PFC signature activity as did YBX1 (**Fig. 5E**). These effects included a comparably high activation of the PFC signature of which YBX1 is a member (PFC Signature 4), raising the possibility of reciprocal activation between YBX1 and this YBX1-driven cohort. Within deep layer neurons, we found that all YBX1-driven PFC TFs are sufficient to activate YBX1 and at least four of the six members of the YBX1-driven cohort (**Fig. 5F, STable 28**). These findings suggest that YBX1 plays an essential role within a small group of TFs that drive each others’ ability to activate PFC fate (**Fig. 5G**).

### Upstream regulators of YBX1-driven PFC determinants

Though YBX1 is essential for expression of most PFC screen targets, we observed three notable exceptions: two established regulators of cortical development (*RORB*, *MECP2*) and the less well-studied *ETV1* (**Fig. 5H**). In our hCO datasets, YBX1 knock-down increased the expression of these PFC-enriched transcription factors. However, in our HPCT dataset, *RORB, MECP2,* and *ETV1* were not among the differentially expressed genes in YBX1-activated cells (**Fig. 5B**) and there was no observable correlation between *YBX1* expression and the levels of either of these three genes (**Fig. 5I**). We therefore hypothesized that RORB, MECP2, and ETV1 represent a group of YBX1-independent PFC determinants.

To determine whether RORB, MECP2, and ETV1 act upstream, in parallel or in opposition to YBX1, we compared the molecular effects of each of these four PFC screening targets in HPCT deep layer neurons. RORB, MECP2, and ETV1 activated PFC signatures in a comparable manner to YBX1 (**Fig. 5J**), and interestingly drove the expression of YBX1 itself, as well as most of the YBX1-driven cohort of PFC determinants (**Fig. 5K**). These results allude to RORB, MECP2, and ETV1 acting as upstream effectors, activating the YBX1-driven cohort and other PFC determinants to specific PFC fate (**Fig. 5L**).

### PFC fate is governed by functionally overlapping cohorts of transcriptional regulators

Synthesizing the findings across our HPCT and hCO datasets alludes to a framework for PFC fate specification, wherein the highly dynamic expression patterns of PFC signatures are coordinated by functionally redundant cohorts of PFC determinants. We further explored this concept by comparing the molecular effects of YBX1-dependent and -independent PFC determinants beyond PFC signatures. Consistent with the highly convergent effects of these determinants on PFC signature activity, we observed an almost complete overlap between the genes and biological processes activated in response to YBX1 and those in response to the YBX1-driven cohort (**SFig. 6C-E, STable 28**). YBX1 and YBX1-independent determinants (*ETV1*, *RORB*, *MECP2*) displayed a similarly comprehensive overlap in downstream gene targets (**SFig. 6F-H, STable 28**). Together, this study proposes a paradigm in which cohorts of intrinsic determinants act as regulatory nodes that collectively orchestrate PFC molecular identity, conferring both the robustness and sensitivity required to integrate extrinsic cues and shape the developing human PFC.

## Discussion

### PFC signatures as tangible readouts of spatiotemporally dynamic areal identities

Our collection of perturbational datasets address long-standing questions of how functionally distinct cortical areas are patterned within the human cerebral cortex. Previous work has focused on dissecting the individual genes that specify these areas in an effort to find their resident master regulators. While this work has revealed important determinants of the visual cortex, including *EMX2*^40^ and *NR2F1*^41^, the full complement of mechanisms that shape the human cortex into as many as 180 cortical areas in the adult^42^ are still unclear.

Area-specific gene programs were largely undetected in bulk transcriptomic profiles of the human cortex^43–45^, and only with granular, developmentally staged bulk transcriptomic comparisons across cortical areas in both human^46^ and non-human primate^47^ did molecular differences between areas become apparent. These studies revealed a “cup-shaped” pattern of areal gene enrichment, in which molecular distinctions between cortical areas diminish during late gestation before re-emerging and amplifying throughout childhood and adolescence^47^, reflecting the complex and dynamic nature of cortical area-specific molecular identity. Subsequent single-cell and spatial transcriptomic profiles of human cortical arealization have corroborated and expanded on this concept^14,48^, including recent work that shows that development segregates in an association vs sensory framework and is highly dynamic^15^.

Taken together, the datasets accrued by the field over the past decades highlight the need for analytical frameworks that capture the dynamic, cell type- and stage-specific nature of cortical area molecular identity across development. Using our meta-module analysis pipeline, we reexamine these seminal datasets to granularly define spatiotemporal and cell type specific signatures. Importantly, our signatures encompass the markers canonically used from bulk studies, but allow us to group these PFC marker genes based on when during development they are most enriched in the PFC and in what cell type. The 18 signatures we present here are valuable for dissecting developmentally dynamic timepoints of PFC development and the strategy could be used for the analysis of other sensory areas in the future.

### *Benchmarked* in vitro *platforms to functionally interrogate cortical arealization*

Functional interrogation of these dynamics is essential to identify drivers of these PFC signatures, but inherently challenging in human development. Cortical organoids provide an opportunity to perform these manipulations: previous studies have identified that area signatures can exist in hCOs^21^ and that extrinsic signals can alter identity^17,22,23,49^. However, given the known limitations of hCOs as models of areal patterning, we sought to systematically identify which PFC signatures, cell type identities, and transcriptional regulatory relationships are preserved relative to HPCT. Our results show that approximately 25% of patterns are preserved between HPCT and hCO, and we notably identify that the cohesion amongst regulators observed in HPCT is much more dissonant in hCO. Importantly, numerous factors are preserved, and this benchmarking allowed us to prioritize the analysis of these candidates, including YBX1.

Despite prior efforts seeking to arealize hCOs, an open question remains as to whether these systems can faithfully recapitulate the dynamic molecular identity of developing cortical areas that underlie conceptual frameworks in the field. For PFC, and possibly association areas specifically, our data suggests we can. Organoids can be used to study the emergence of gene programs related to association area identities, though as we show, the fidelity to primary tissue is for a subset of transcriptional regulators, PFC signatures, and cell types. Future studies linking these isolated molecular programs to structures of other cortical areas will be essential to more fully understand the emergence and crosstalk between cortical areas.

### Novel cohorts of PFC determinants

Across our pooled and targeted CRISPR perturbations in HPCT and hCO models, YBX1 consistently emerged as a necessary and sufficient activator of both PFC signatures and PFC determinants. This multifunctional protein has well-known regulatory functions in a broad range of essential cellular processes^50–52^ and nearly every aspect of transcriptional and translational regulation^53^. YBX1 activity in the context of cancer^34–36^ and neuronal function^37,38,54^ has been well described, and recent reports in the mouse cortex point to an essential role for YBX1 in maintaining radial glia proliferation^26^. However, the roles of YBX1 in human cortical development, and specifically in PFC specification, have not been previously examined.

Our HPCT and hCO platforms enable us to test the neurodevelopmental roles of YBX1 in the context of human cortical tissues. Consistent with the models suggested by analogous studies in the mouse cortex, YBX1 knock-down experiments in hCOs resulted in an increase in deep layer neuronal fates^26^. However, activation of YBX1 in hCOs or HPCTs did not result in a corresponding increase in radial glia proliferation, and in fact, resulted in slight increases in deep layer neuronal fates. These findings suggest that, while the loss of YBX1 consistently disrupts the cortical progenitor pool, as-yet unknown regulatory mechanisms are in place to prevent elevated YBX1 levels from driving progenitor expansion in the developing human cortex.

YBX1 protein levels have been reported to be under tight control^35^, with studies demonstrating its ability to bind to its own mRNA^37,55^ and reduce its own translation^55^. Our screening datasets reveal nine activators of YBX1, including MECP2, which has been shown to physically interact with YBX1 and act in concert to regulate mRNA splicing^38^. Such regulatory feedback loops may underlie the intriguing, switch-like responses we observed for the six YBX1-sensitive PFC signatures identified in both HPCT and hCO screens. More broadly, these functional relationships between YBX1 and other PFC-enriched TFs may represent fail-safe mechanisms that ensure robust PFC development. These feedback relationships amongst PFC determinants mirror the recently described autoregulatory loops between retinoic acid, its synthesizing enzyme ALDH1A3, and MEIS2 (a target evaluated in our hCO screen) to maintain the boundary between the PFC and motor areas^56^. Alternatively, subtle differences in the ability of these PFC activator cohorts can enable the PFC to react to subtle, extrinsic cues, including retinoic acid signaling^17^.

### The link between area-specific molecular signatures and unique cognitive functions

Our findings support frameworks for PFC specification that move beyond constant, static marker genes, and towards a model of distinct TF cohorts that work in tandem to activate and sustain the correct molecular signatures at the appropriate developmental timepoints. These concepts mirror recent work that describes antagonistic relationships between the molecular programs of association areas like the PFC and sensorimotor areas like the V1^15^. As stated above, the mechanisms in our perturbational datasets may also interact with the recently described postmitotic feedback loops that strengthen the retinoic acid-induced boundaries of the PFC15. While retinoic acid regulators were enriched in one of our PFC signatures (signature 9), our screens were unable to identify TFs that could strongly activate this gene program. Signature 9 may highlight a limitation in our *in vitro* systems to model the extrinsic determinants of areal fate, and more broadly, the consequences of these molecular determinants on neuronal circuitry and cognitive function.

However, these recent studies^15,56^ have demonstrated the essential role of areal determinants on connectivity and cognition. In mouse models, the depletion of molecular determinants of association and sensorimotor areas resulted in changes in axon guidance, the absence of thalamocortical afferents, and altered projections to the PFC^15^. A separate study additionally found that *Meis2* depletion not only shrank the frontal cortical areas in favor of expanded motor cortex, but also led to disrupted thalamocortical projections that translated to deficits in working and spatial memory^56^. Our *in vitro* perturbation screens complement this work, enabling high-throughput, genetically tractable mechanisms to isolate the intrinsic mechanisms by which gene networks initiate the patterning of human cortical areas.

## Acknowledgements

We would like to thank the members of the Bhaduri Lab for their insightful advice and comments on the study. We would also like to thank the members of the Broad Stem Cell Research Center Flow Cytometry Core for their help in isolating cells for this project; C. Julian (UCSF Institute for Human Genetics Genomics Services Core) and S. Feng (UCLA Broad Stem Cell Research Center Sequencing Core). The work performed in the manuscript was generously funded by R00NS111731 from the NIH (NINDS), R01MH132689 from the NIH (NIMH), the Young Investigator Award from the Brain & Behavior Research Foundation, the Alfred P. Sloan Foundation, the Rose Hills Foundation, the Klingenstein-Simons Fellowship from the Esther A. & Joseph Klingenstein Fund and the Simons Foundation, the NIH BRAIN Initiative Cell Atlas Network (UM1MH130991), and the Ablon Scholar Award (to A.B.). Additional funding was provided to P.R.N. (UCLA Zamenhof Scholarship, UCLA Eli and Edythe Broad Center of Regenerative Medicine and Stem Cell Research Training Program), J.A.S. (Eugene V. Cota-Robles Award), J.M. and A.M. (UCLA Eli and Edythe Broad Center of Regenerative Medicine and Stem Cell Research Training Program), and to M.H. (NIMH BRAIN, NIMH RF1MH132662, NHGRI U24HG002371 and CIRM DISC0-14514 (with A.B.)). This material is based upon work supported by the National Science Foundation Graduate Research Fellowship Program under Grant No. (DGE-2034835) to J.A.S. Any opinions, findings, and conclusions or recommendations expressed in this material are those of the author(s) and do not necessarily reflect the views of the National Science Foundation.

## Author Contributions

The study was conceptualized by PRN and AB. Experimental designs were made by PRN, DCJ, and AB. Experiments were performed by PRN, DCJ, VG, and SW. JAS, JM, and AM assisted with experiments, HPCT processing, and sequencing. Formal analysis was conducted by PRN, with support from DCJ. Data visualization was performed by PRN, BW and MH. Data interpretation was performed by PRN, DCJ and AB. Project supervision, administration, and funding acquisition were provided by AB. The manuscript was written by PRN and AB with input and edits from all authors.

## Data Availability

Our perturbational dataset of HPCT pooled CRISPR screens is browsable in the UCSC Genome Browser (https://cells-test.gi.ucsc.edu/?ds=dev-hpct-pfc-perturb), in which our dataset is also available as a Seurat object for download.

## Methods

### Curation of PFC and V1 marker genes from bulk transcriptomic datasets

Genes with the greatest correlation between gene expression and position along the rostrocaudal axis were identified from an existing, bulk transcriptomic profile of the developing human cortex^16^. The subsequent initial bulk PFC and V1 marker sets were then validated in a single-cell transcriptomic dataset of developing cortical areas^14^. Using our module activity scoring^24^, we calculated the average gene expression of these marker sets. Briefly, the activity score for each gene set was calculated in each cell by summing the normalized counts per million (CPM) for all of the genes in the set, then dividing by the total number of set genes. We then measured the correlation of area marker set activity vs the expression of each individual marker gene in this dataset, isolating the marker genes which demonstrated expression levels that were highly correlated with marker set activity. These analyses narrowed our marker sets to the 32 bulk PFC marker genes and 10 bulk V1 marker genes shown in Fig. 1A.

### PFC signature generation

PFC signatures were generated by applying our meta-module pipeline^24^ to a single-cell RNA-seq profile of the developing human PFC and V1^14^ spanning early (GW 14–17, n = 3), mid (GW 18–20, n = 6), and late (GW 22, 25, n = 4) developmental stages. The input for meta-module generation consisted of differentially expressed genes previously calculated for each individual sample. Specifically, PFC- and V1-enriched genes were determined within individual cell type populations and scored using a gene score metric that quantifies the enrichment and specificity of a gene to its assigned area^14,24^. Marker genes were then filtered to the 90th percentile of gene score within each developmental stage-area-cell type combination. The resulting table of marker genes and their calculated gene scores across combinations of developmental stage, area, and cell type was used to generate a distance matrix. Hierarchical clustering of this matrix was used to group genes based on their specificity and enrichment to the PFC vs V1, across cell types and developmental stages.

The activity of the resulting 21 modules was then evaluated in PFC and V1 samples from our meta-atlas of developing human primary cortical tissues^24^. Any module with a statistically significant (two-sided Wilcoxon test), greater than 25% increase in activity in PFC versus V1 was designated as a PFC signature, resulting in 18 PFC signatures encompassing 545 genes (**STable 3**). Gene ontology term enrichment analysis was conducted on each PFC signature using the enrichR R package (v3.2) surveying signaling pathway databases (WikiPathway 2021 Human, KEGG 2021 Human, Elsevier Pathway Collection), transcriptional regulatory collections (ChEA 2016, ENCODE and ChEA Consensus TFs from ChIP, TF Perturbations Followed by Expression, TRRUST Transcription Factors 2019), and gene ontology sets (GO Biological Process 2021, GO Molecular Function 2021, GO Cellular Component 2021) to annotate the biological processes represented by each signature (**STable 4-5**).

### Identification and curation of PFC screen target genes

PFC screen target genes were selected from PFC- and V1-enriched marker genes identified in the same single-cell RNA-seq profile used to originate our PFC signatures^14^. To be considered for inclusion, a gene was required to: (1) be identified as an areal marker in at least one of the following cell type contexts: exclusively in radial glia, indicative of upregulation at the initiation of areal patterning; or across radial glia, IPCs, and neurons, indicative of upregulation throughout the entire cell lineage of a given area; and (2) be a transcription factor, as defined by the ChEA Transcription Factor Dataset, indicative of a likely gene regulatory role in areal patterning.

Candidate genes passing these criteria were then evaluated for their expression in human cortical organoids (hCOs) relative to primary tissue ^14,21^. Genes were prioritized for inclusion if their average log2 fold-change of primary tissue versus organoid expression was greater than or equal to −1, indicating that their expression is not substantially higher in organoids than in primary tissue and that ectopic upregulation in organoids would therefore be expected to have a meaningful effect on areal fate. Genes for which organoid expression substantially exceeded primary tissue expression were excluded. Genes that were enriched as areal markers across all cell types of both PFC and V1, or that marked only radial glia of the target area but all cell types of the opposing area, were also excluded given the difficulty in interpreting area-specific effects. Genes enriched across all cell types of the target area but only in radial glia of the opposing area were retained, as their broad marker status in the target area is more consistent with a role in area-specific patterning. These criteria collectively yielded 35 PFC screen target genes (**STable 6**).

Guide RNA (gRNA) sequences for all 35 PFC screen target genes were selected from the CRISPRa v2 collection^57^, in which gRNA sequences were designed using a machine learning algorithm trained on prior CRISPRi and CRISPRa screening data. For each target gene, the top 7 sgRNA sequences ranked by predicted activity were included. Where a target gene possessed both a primary and secondary transcription start site meeting the FANTOM consortium threshold, sgRNAs were designed against both sites; in cases where these sites were separated by approximately 1 kb, two independent sets of 7 sgRNAs were included for that gene, yielding 294 targeting gRNAs in the library (**STable 7**). Non-targeting control gRNAs were selected from the h5 (Gene Expression) sublibrary of the hCRISPRav2 non-targeting collection and represented at 21-fold greater abundance relative to any individual targeting gRNA, following the approach of Replogle et al., 2021^33^ and Datlinger et al., 2017^58^, to ensure comprehensive representation of all major cell types in the neuronal lineage among non-targeting control-expressing cells.

### Generation of lentiviral constructs

The dCas9-VP64 activator construct used in all CRISPRa experiments was expressed from pLV hUbC-VP64 dCas9 VP64-T2A-GFP (Addgene #59791).

The pooled gRNA library used for CRISPR activation screens was cloned into the pU6-sgRNACROP-mCherry backbone (Addgene #271288) using a pooled, amplification-free Gibson Assembly approach based on Datlinger et al., 2017^58^. Briefly, the pU6-sgRNACROP-mCherry vector was linearized by overnight digestion with BstXI and BlpI at 37°C, and the linearized vector was gel-purified. Pooled oligonucleotide libraries (oPools, IDT) encoding all gRNA sequences and appropriate overhanging sequences were resuspended to 100 µM per oligo in EB buffer and diluted to a working concentration of 100 nM. Gibson Assembly reactions were set up using NEBuilder HiFi DNA Assembly Master Mix (NEB) at a molar ratio of 11 fmol vector : 200 fmol insert in a 20 µL reaction and incubated for 1 hr at 50°C. Assembled products were electroporated into Lucigen Endura competent cells (25 µL cells + 10 µL Gibson Assembly product) using the following parameters: 25 µF capacitance, 200 ohms, 1.5 kV. Cells were recovered in Lucigen Endura Recovery Media at 30°C for 1 hr with shaking, then plated onto LB + 100 µg/mL ampicillin plates and incubated at 30°C for 20 hours. Colonies were scraped from plates in LB, pelleted at 5,000 × g for 15 min at 4°C, and plasmid libraries were prepared using a Macherey-Nagel Maxiprep EF kit (#740424). Library representation was confirmed by MiSeq sequencing following Replogle et al., 2021^33^, using PCR amplification with staggered P5 primers annealing to the mU6 promoter (mU6_F1: 5′-cagcacaaaaggaaactcacc-3′) and indexed P7 primers annealing to the CROP-seq scaffold (CROP_R2: 5′-caagttgataacggactagcc-3′), 12 cycles of amplification using Q5 polymerase, and SPRI bead cleanup at a 0.9X ratio (Beckman Coulter, B23318).

For YBX1 knock-down experiments, YBX1-targeting shRNA constructs were generated by annealing and ligating oligos encoding the shRNA sequence (sense strand: 5′-CCAGTTCAAGGCAGTAAATAT-3′, IDT) into PacI/EcoRI-linearized pLKO.3G-EGFP (Addgene #14748) using T4 DNA ligase (16°C, overnight).

### Lentiviral production

For large-scale lentiviral production of dCas9-VP64, HEK-293T cells (Takara, Cat. # 632180) were seeded at 90% confluency across 22 × 10-cm plates and transfected with 5.9 µg pMD2.G (Addgene #12259), 9.4 µg psPAX2 (Addgene #12260), and 10.2 µg pLV hUbC-VP64 dCas9 VP64-T2A-GFP per plate using Lipofectamine 2000 (Cat. # 11668019, Thermo Fisher) in Opti-MEM Reduced Serum Media. Medium was replaced after 24 hours with complete HEK-293T culture media (10% FBS, 1X penicillin-streptomycin in high-glucose DMEM supplemented with GlutaMAX and pyruvate, Cat. # 10569010, Thermo Fisher). Supernatants were collected at 2 and 3 days post-transfection, passed through a 0.45-µm filter, concentrated with Lenti-X Concentrator (Cat. # 631232, Takara) by overnight incubation (∼18 hours) at 4°C. Supernatants were then pelleted by centrifugation at 1,500 × g for 1 hour at 4°C, and pellets were resuspended in 1/150th of the original volume in cold HPCT Media or Organoid Media 1, aliquoted, and stored at −80°C.

Lentiviral supernatants for pooled gRNA libraries and YBX1 shRNAs were generated as described above for dCas9-VP64, using 10-cm plate per lentivirus and a final resuspension in PBS. Plasmid transfection amounts were 1.3 µg pMD2.G, 5.8 µg psPAX2, and 5.8 µg pU6-sgRNACROP-mCherry for gRNA libraries. For YBX1 shRNA, 3 µg pMD2.G, 9 µg psPAX2, and 12 µg pLKO.3G-YBX1 shRNA-EGFP was used.

### Pooled CRISPR screening in HPCT

HPCT samples were obtained and processed as approved by Ronald Reagan UCLA Medical Center Translational Pathology Core Laboratory. Cultures were derived from four samples spanning the first trimester (GW 7 and GW 10) and second trimester (GW 14 and GW 15) of gestation and dissociated and plated as previously described^31^. For each screen, dissociated cortical tissue was plated as a monolayer on poly-D-lysine-coated 12-well plates (Thermo, A3890401; 50 µg/mL, 1 hr coating) at approximately 1 million cells per well. HPCT cells were cultured in DMEM/F12-GlutaMAX (Cat. # 10565-018, Life Technologies) supplemented with 1X B-27 (Cat. # 17504044, Thermo Fisher), 1X N2 (Cat. # 17502-048, Life Technologies), 1X sodium pyruvate (Cat. # 11360070, Life Technologies), and 1X penicillin-streptomycin (HPCT Media). Twenty-four hours after plating, cells were lentivirally infected with dCas9-VP64-expressing supernatants (5% v/v) and PFC screen gRNA library-expressing supernatants (0.1% v/v). Media was changed approximately 20 hours after infection, and subsequently every other day.

Following 8–10 days of culture, cells were dissociated for FACS. Wells were washed twice with warm PBS/BSA (0.5% BSA in Ca/Mg-free PBS), then incubated with papain (Cat. # LK003176, Worthington) in dissociation media [HBSS supplemented with HEPES, kynurenic acid, MgClL, and glucose, pH 7.35] containing 10 µM Rock inhibitor Y-27632 (Cat. # 72304, Stem Cell Technologies), 5 µg/mL Actinomycin D (Cat. # A9415, Sigma), and 10 µg/mL Anisomycin (Cat. # A5862, Sigma) for approximately 1-6 min at 37°C. Papain was quenched by addition and immediate removal of cold ovomucoid inhibitor solution, and cells were resuspended in cold DNAse solution in dissociation media, triturated 10 times with a P1000 pipette, and transferred to a 50 mL conical tube. Samples were diluted to 5 mL per well with additional DNAse solution and pelleted at 300 × g for 5 min at 4°C. Cell pellets were resuspended in 500 µL PBS/BSA, passed through a pre-wetted 40 µm cell strainer, and treated with DAPI (1:1000). GFP+/mCherry+ (dCas9a+; sgRNA+) cells were sorted on a FACSAria H or FACSAria (BD Biosciences) with 100 µm nozzles at the BSCRC FACS core facility. GFP+/mCherry+ cells were pooled across wells and captured using the 10X Genomics 3′ v3.1 kit (Dual Index) (10X Genomics, CG000315 Rev F) or 10X Genomics High-throughput 3′ v3.1 kit (Dual Index) (10X Genomics, CG000417 Rev D) according to manufacturer instructions. Sequencing libraries were generated using 11-12 cycles for cDNA amplification, and 11-12 cycles for gene expression library amplification depending on input DNA amount. Libraries were quantified on an Agilent 2100 Bioanalyzer and sequenced on a NovaSeq S1 or S2 flow cell.

### Standard stem cell maintenance and cortical organoid generation

Human embryonic stem cell lines UCLA1 hESC and UCLA6 hESC, and the induced pluripotent stem cell line KOLF iPSC, were authenticated at source. Stem cells were thawed in mTeSR+ media (Cat. # 100-0276, Stem Cell Technologies) and plated onto six-well plates coated with growth factor-reduced Matrigel (Cat. # 354230, Corning). Cells were treated overnight with 10 μM Rock inhibitor Y-27632 (Cat. # 72304, Stem Cell Technologies) to attenuate cell death. Stem cells were subsequently maintained in mTeSR+, with media changed every 1–3 days. Cells were passaged at approximately 70% confluency, at which time cells were washed with PBS and incubated in ReleSR (Cat. # 05872, Stem Cell Technologies) for 1 min at room temperature. The ReleSR was aspirated, and cells were incubated at 37°C for 5 min, after which mTeSR+ was added to the well and cells were manually lifted with cell lifters (Cat. # 08-100-240, Fisher) and expanded onto Matrigel-coated six-well plates.

In general, cortical organoids were generated based on protocols described in Kadoshima et al., 2013^18^ and Velasco et al., 2019^19^. Stem cells were washed with PBS and dissociated into single-cell suspensions with a 5-min, 37°C incubation in Accutase (Cat. # A6964, Millipore Sigma). Cells were scraped and pelleted for 5 min at 300 × g, then immediately resuspended in Organoid Media 1 [GMEM (Cat. # 11710-035, Life Technologies) supplemented with 1X penicillin-streptomycin or primocin, 15% KnockOut Serum (Cat. # 10828-028, Life Technologies), 1X NEAA (Cat. # 11140050, Life Technologies), 1X sodium pyruvate (Cat. # 11360070, Life Technologies), 100 μM β-mercaptoethanol (Cat. # 21985023, Life Technologies), 20 μM Rock inhibitor Y-27632, 10 μM SB431542 (Cat. # 1614, Tocris), and 3 μM IWR1-e (Cat. # 13659, Cayman Chemicals)]. Resuspended cells were transferred to 96-well v-bottom low-adhesion plates (Cat. # MS-9096VZ, S-Bio) at 10,000–13,000 cells per well.

Throughout cell culture, media was changed every 2–3 days. For the first 6 days, organoids were treated with 20 μM Rock inhibitor Y-27632. After 18 days, organoids were transferred to six-well ultra-low adhesion plates (Cat. # 07-200-601, Corning) and placed onto an orbital shaker rotating at 100 rpm. At this time, organoids were moved to Organoid Media 2 [DMEM/F12-GlutaMAX media (Cat. # 10565-018, Life Technologies) supplemented with 1X N2 (Cat. # 17502-048, Life Technologies), 1X CD Lipid Concentrate (Cat. # 11905-031, Life Technologies), and 1X penicillin-streptomycin or primocin]. Beginning at day 35, organoid culture media was supplemented with 10% heat-inactivated FBS (Cat. # 10082147, Life Technologies), 5 μg/mL heparin (Cat. # H3149, Sigma), and 0.5% Matrigel (Media 3).

### Pooled CRISPR screening in hCOs

Pooled gRNA library was introduced into UCLA1 hESC stem cells through lentiviral infection, at least two passages post-thaw. One ∼70% confluent six-well well of stem cells was passaged and expanded into three wells, with each resulting well containing 10 µM Rock inhibitor Y-27632, 10 µg/mL polybrene (Cat. # TR-1003-G, Millipore Sigma), and gRNA library lentiviral supernatant at a 1:100 dilution in mTeSR+ media. Media was changed approximately 18 hours after infection.

Infected stem cells were then used to generate cortical organoids as described above. At day 17, dCas9-VP64 was introduced with an adaptation of the chimeroid approach^24,39^. Organoids were dissociated into single-cell suspensions via 30-min incubation at 37°C in papain and DNAse in EBSS (Cat. # LK003150, Worthington), shaking vigorously every 5 min. Samples were triturated 10 times with a P1000 pipette, after which ovomucoid inhibitor solution was added to quench papain activity, and cells were pelleted with a 5-min spin at 300 × g at 4°C. Cell pellets were resuspended in Organoid Media 1 and transferred to 96-well v-bottom low-adhesion plates at 30,000 cells per well, simultaneously incubated with dCas9a lentiviral supernatants (pLV hUbC-VP64 dCas9 VP64-T2A-GFP) at a 10% v/v dilution.

Re-aggregated cells were incubated in lentivirus for 18 – 24 hrs before media was changed to baseline Organoid Media 1. At day 19, media was transitioned to Organoid Media 2, and at day 24, organoids were transferred to six-well ultra-low adhesion plates on an orbital shaker at 100 rpm, with media supplementation progressing as described for standard organoid generation.

At 8 weeks of culture, organoids were quality-controlled and non-cortical regions were dissected. Organoids were then dissociated via incubation in papain and DNAse in EBSS (Cat. # LK003150, Worthington) at 37°C, shaking vigorously at 5-min intervals for a minimum of 30 min, followed by trituration with a P200 pipette. Cells were pelleted at 300 × g for 5 min at 4°C, resuspended in FACS buffer (PBS + 2% FBS), passed through a 35-µm cell strainer, and treated with DAPI (1:1000). Cells were sorted on a FACSAria (BD Biosciences) with a 100 µm nozzle at the BSCRC FACS core facility to isolate mCherry+;GFP+ cells. Sorted cells were captured using the 10X Genomics High-throughput 3′ v3.1 kit (Dual Index) (10X Genomics, CG000417 Rev D) according to manufacturer instructions. Sequencing libraries were generated using 11 cycles for cDNA amplification, and 12 cycles for gene expression library amplification. Libraries were quantified on an Agilent 2100 Bioanalyzer and sequenced on a NovaSeq S1 or S2 flow cell.

### YBX1 knock-down in hCOs

Cortical organoids were generated from each of UCLA1 hESC, UCLA6 hESC, and KOLF iPSC lines as described above. At day 19, organoids from all three lines were dissociated and re-aggregated together in the presence of YBX1 shRNA lentivirus using a chimeroid-based approach^24,39^, incorporating improvements from the hCO pooled CRISPR screening. For each line, organoids were transferred to a 5 mL tube, excess media was removed, and papain and DNAse solution in EBSS (Cat. # LK003150, Worthington) was added. Samples were incubated for 10 min at 37°C, shaken by hard inversion 30 times, incubated for an additional 10 min, and triturated 20 times with a P200 pipette until near-single-cell suspension was achieved. Cells were pelleted at 300 × g for 5 min at room temperature and resuspended in Organoid Media 1. Single-cell suspensions from all three lines were combined in equal proportions and transferred to 96-well v-bottom low-adhesion plates at 100 µL per well, simultaneously incubated with YBX1 shRNA lentiviral supernatants at a 1% v/v dilution.

At 18–20 hours post-infection, 100 µL of media was replaced with Organoid Media 2. Media was changed again the following day. At day 23, organoids were transferred to six-well ultra-low adhesion plates on an orbital shaker at 100 rpm, with media supplementation progressing as described for standard organoid generation.

Following 9 weeks of culture, the efficacy of YBX1 shRNA constructs was validated in EGFP+ (YBX1 shRNA+) and EGFP− (YBX1 shRNA−) cells were isolated by FACS from the same organoids. Samples were prepped for FACS as described for the pooled CRISPR screening in hCOs, and cells were sorted on a FACSAria (BD Biosciences) with a 100 µm nozzle at the BSCRC FACS core facility. RNA was extracted using the RNeasy Plus Mini Kit (Qiagen), and cDNA was synthesized from 160 ng RNA per reaction using the SuperScript IV VILO Master Mix (Cat. # 11756050, Thermo Fisher). Quantitative PCR was performed in triplicate 10 µL reactions using SYBR Green PCR Master Mix (Cat. # 4368708, Thermo Fisher) on a QuantStudio 5 (384-well block) using the Comparative-Ct (ΔΔCT) method, with GAPDH as the housekeeping gene. Primer sequences used: YBX1 (forward: 5′-TGGGCGTCGACCACAGTAT-3′; reverse: 5′-CAGCACCCTCCATCACTTCTC-3′) and GAPDH (forward: 5’-GGAGCGAGATCCCTCCAAAAT-3’; reverse: 5’-GGCTGTTGTCATACTTCTCATGG-3’).

A week later (10 weeks of culture), EGFP+ (YBX1 shRNA+) and EGFP− (YBX1 shRNA−) cells were isolated from organoids for single-nuclei multiomic analysis. FACS was performed as described above, but sorted cells were processed for nuclei isolation per the 10X Nuclei Isolation Protocol (CG000365), using a 3-min lysis buffer incubation and three 1-mL Wash Buffer washes. Multiomic libraries were generated using the Chromium Next GEM Single Cell Multiome ATAC + Gene Expression Kit (10X Genomics, CG000338 Rev F) according to manufacturer instructions, using 7-8 cycles for ATAC library amplification, 6-7 cycles for cDNA amplification, and 10-14 cycles for gene expression library amplification depending on input DNA amount. Libraries were quantified on an Agilent 2100 Bioanalyzer and sequenced on a NovaSeq X Plus flow cell.

### Evaluation of cortical organoid patterning

After 5 weeks of culture, organoids and chimeroids were fixed for 45 min in 4% paraformaldehyde, washed with 1X PBS, and rehydrated with an incubation in 30% sucrose in PBS at 4°C overnight. Organoids were embedded into cryomolds in a 1:1 ratio of 30% sucrose in PBS and OCT (Cat. # 14-373-65, Tissue-Tek), placed flat on dry ice for 10 min, and stored at −80°C. Sections were cut on a cryostat at 16 - 20 µm and mounted onto treated glass slides. After a 5-minute PBS wash, sections were incubated with boiling citrate-based antigen retrieval solution (Cat No. H-3300-250; Vector Laboratories) for 20 minutes, then washed with PBS and permeabilized with 0.1% Triton in PBS for 10 minutes. Slides were then incubated in blocking buffer (3% BSA, 5% donkey serum, and 0.05% Tween-20) for 30 min, then incubated overnight at 4°C with primary antibodies diluted in blocking buffer. Following washes with 0.05% Tween-20 in PBS, slides were incubated in the dark at room temperature for 2 hours with 1:1000 DAPI and 1:500 AlexaFluor secondary antibodies (Thermo Fisher) diluted in blocking buffer. Primary antibodies used: mCherry (1:500, Cat. #43590, Cell Signaling Technologies), GFP (1:500, Cat. # GFP-1020, Aves Labs), SOX2 (1:500, Cat. # sc-365823, Santa Cruz Biotechnology or Cat. # AB97959, Abcam), CTIP2 (1:500, Cat. # ab18465, Abcam). Stained organoids were imaged on an EVOS M5000 (Thermo Fisher) with 4X and 10X objectives, and fluorescence intensities later adjusted using FIJI software (RRID:SCR_002285). Image acquisition and adjustment parameters were left constant for all samples within the same staining panel.

### Generation of gRNA-enriched side libraries

For all screens, gRNA-enriched side libraries were generated from GEX cDNA to enable single-cell gRNA detection, following Replogle et al., 2021^33^. Briefly, one-sixth of the amplified cDNA from each GEX capture was used as input per 100 µL PCR reaction for amplification of the CROP-seq scaffold, using KAPA HiFi HotStart ReadyMix for 25 cycles with the following primers:

TruSeqR2-CROP_F3_1: 5′-

GTGACTGGAGTTCAGACGTGTGCTCTTCCGATCTCTTGGAGAACCACCTTGTTG-3′

prtlTruSeqR1_1:

5′-ACACTCTTTCCCTACACGACG-3′

For samples captured using the 10X Genomics High-throughput 3′ v3.1 kit, two technical replicate PCR reactions were run per sample to account for the larger cDNA input from the high-throughput capture. PCR products were purified with a 0.7X SPRI bead cleanup (Beckmann Coulter, B23318) and eluted in EB buffer. Sample index PCR was performed for 5 cycles following the 10X Genomics 3′ v3.1 library construction protocol, and libraries were purified by double-sided SPRI (0.5X then 0.3X) to yield a final product of approximately 630 bp. Libraries were quantified on an Agilent 2100 Bioanalyzer, and sequenced as per manufacturer recommendation on a NovaSeq X Plus flow cell. Side library FASTQ files were aligned to a custom reference containing the full CROP-seq scaffold sequences for all 315 gRNAs in the PFC screen library using CellRanger v6.1.1, and raw (unfiltered) barcode matrix outputs were used as input to the context-aware gRNA detection pipeline to maximize the recovery of perturbed cells.

### Context-aware, single-cell gRNA detection pipeline

To rigorously identify cells harboring effective perturbations, we developed a context-aware, single-cell gRNA detection pipeline. The pipeline proceeds in three steps, applied independently for each PFC screen target gene and separately for each of the HPCT and hCO pooled CRISPR screen datasets (Supplementary Tables 7–8).

First, valid non-targeting (NT) gRNAs were identified for each library target gene. NT gRNAs were evaluated in a query population of cells expressing only one NT gRNA. For each NT gRNA, the expression of each screen target gene was compared between query cells and a negative control population of all cells expressing no targeting gRNAs, using a two-sided Wilcoxon rank sum test. A NT gRNA was designated as valid for a given target gene if it did not significantly alter that gene’s expression (p > 0.05). Valid NT gRNAs were therefore defined on a per-target-gene basis, such that the same NT gRNA could be valid for one target gene but invalid for another.

Second, valid targeting gRNAs were identified for each library target gene. For each targeting gRNA, the expression of its target gene was compared between cells expressing that gRNA and a negative control population of cells expressing only the valid NT gRNAs for that target gene and no targeting gRNAs, using a one-sided Wilcoxon rank sum test (alternative = ‘greater’; p < 0.05). A targeting gRNA was designated as valid if it significantly upregulated its target gene’s expression relative to this gene-specific negative control population. Target genes for which no valid targeting gRNAs were identified were excluded from downstream analyses.

Third, individual cells were classified as perturbed or unperturbed for each target gene. For each target gene, the counts of all valid targeting gRNAs per cell were summed, and a per-cell gRNA count threshold was determined using a Gaussian-based local minima approach applied to the density of log2 gRNA counts^33^. Briefly, a kernel density estimate was computed on log2 gRNA counts, and the minimum of the lowest local minimum of the density was used as the count threshold. In cases where fewer than three unique log2 count values were present, a binary threshold of 0 was applied. Cells exceeding this threshold were classified as expressing a valid targeting gRNA for that target gene and designated as perturbed. Cells were assigned a perturbation status listing all PFC screen targets for which they were classified as perturbed, enabling classification into unperturbed, singly perturbed, and multiply perturbed populations.

### Standard single-cell/nuclei RNA-seq data processing and quality control

FASTQ files from HPCT and hCO pooled CRISPR screen experiments were aligned to a custom reference containing the human genome (GRCh38-2020-A), genomes of four common mycoplasma species, a dCas9a-GFP sequence, and an mCherry-CROP sequence using CellRanger v6.1.1, with intronic reads included (--include-introns=TRUE). For the YBX1 knock-down multiomic experiment, FASTQ files were aligned to a custom reference containing the human genome (GRCh38-2020-A), genomes of four common mycoplasma species, and the EGFP reporter sequence from pLKO pLKO.3G-EGFP using CellRanger v8, with --chemistry=ARC-v1 and --include-introns=true. For all datasets, filtered barcode matrices were converted to Seurat objects using the Read10X function with min.cells = 3 (v5). Prior to merging, putative doublets were identified and removed per well or per sample using DoubletFinder (v2.0.4^59^), with pN = 0.25, pK selected by BCmetric maximization, and nExp calculated from empirically estimated per-sample multiplet rates. Following doublet removal, samples were merged and quality control was applied by omitting cells with fewer than 1,000 genes detected and more than 5% of UMIs mapping to mitochondrial genes. For the YBX1 knock-down GEX data, ambient RNA contamination was additionally removed using CellBender (remove-background, --cuda, v0.3.2^60^) prior to DoubletFinder, and the QC threshold was set at fewer than 500 genes detected. Data were normalized using LogNormalize (scale.factor = 10,000).

Three merged Seurat objects were generated, each containing (1) all samples from the HPCT screens; (2) all sorted cells from the hCO screen; or (3) all sorted cells from the YBX1 knockdown experiment. For each merged Seurat object, cells were clustered using conventional Seurat pipelines to annotate cell types. Data were normalized using LogNormalize (scale.factor = 10,000), variable features were identified using FindVariableFeatures (vst, nfeatures = 2,000), and data were scaled prior to principal component analysis (RunPCA). Significant principal components were identified using methods described in Shekhar et al., 2016^61^, and used to run a graph-based clustering approach using the FindNeighbors and FindClusters functions, and cells were visualized using RunUMAP. Cluster markers were identified using FindAllMarkers, and gene scores were calculated as described above^24^. Cell type identities were assigned based on canonical marker gene expression. The cortical identity of HPCT samples was additionally validated using the label transfer pipeline in Seurat (FindTransferAnchors and MapQuery), which was used to transfer cell type labels from a first-trimester^32^ and second-trimester cortical reference^14^.

### Gene program activity scoring and comparison

Developmental meta-module and PFC signature activities were calculated per cell using our module activity score^24^, as described above. To evaluate the effects of each perturbation on gene program activity, the percent change in average gene program activity between perturbed and unperturbed cells was calculated for each screen target gene and each gene program, within each cell type as indicated. Significant changes in activity were identified using two-sided Wilcoxon rank sum tests (p < 0.05). Analogous analyses were conducted to compare gene expression and chromatin accessibility of the indicated gene programs as a result of YBX1 knock-down. For PFC signature calculations, PFC screen targets (HPCT, hCO screen) and YBX1 (YBX1 knock-down) were excluded from gene lists prior to scoring to prevent the perturbed gene from influencing gene program activity estimates.

The Spearman correlation between each PFC screen target gene’s expression and each PFC signature’s activity was calculated within unperturbed and singly perturbed cells, across all cell types and replicate experiments, using the ggpubr R package.

### Differential gene expression and gene ontology analysis

Differentially expressed genes (DEGs) between perturbation conditions were identified using the FindMarkers function (Seurat; p < 0.05, two-sided Wilcoxon rank sum test). Gene ontology term enrichment analysis of DEG sets was conducted using the enrichR R package (v3.2) surveying signaling pathway databases (WikiPathway 2021 Human, KEGG 2021 Human, Elsevier Pathway Collection), transcriptional regulatory collections (ChEA 2016, ENCODE and ChEA Consensus TFs from ChIP, TF Perturbations Followed by Expression, TRRUST Transcription Factors 2019), and gene ontology sets (GO Biological Process 2021, GO Molecular Function 2021, GO Cellular Component 2021).

### Cell line demultiplexing

Cell lines in chimeroid YBX1 knock-down datasets were distinguished as described previously^24^. Briefly, SNP-based genomic probabilities were generated for each of the three stem cell lines (UCLA1 hESC, UCLA6 hESC, and KOLF iPSC) using a Genomic Diversity Array. Biallelic variants were filtered using bcftools and used as reference genotypes to demultiplex individual cell lines using Vireo (run via Demuxafy). SNP pileup was generated from sorted BAM files using cellsnp-lite (--minMAF 0.03, --minCOUNT 5, --UMItag None). Cells were assigned to donor cell line identities using the best_singlet results from the Vireo output.

### Chromatin accessibility analysis

For the YBX1 knock-down multiomic dataset, a unified peak set was generated by merging ATAC peaks called by CellRanger across YBX1 knock-down and unperturbed samples using GenomicRanges::reduce, filtering to retain peaks between 20 bp and 10,000 bp in width and excluding peaks on non-standard chromosomes. Chromatin assays were created using the counts from this peak set (Signac v1.14^62^; min.cells = 10, min.features = 200), with gene annotations extracted from EnsDb.Hsapiens.v86 (hg38). Quality control metrics (nucleosome signal score, TSS enrichment score, fraction of counts in blacklist regions, fraction of reads in peaks) were calculated using built-in Signac functions, and cells were retained if they met all of the following thresholds: nCount_ATAC > 1,000 and < 150,000; FRiP > 0.35; blacklist fraction < 0.05; nucleosome signal < 2; TSS enrichment > 1. The GEX and ATAC assays were intersected to retain only cells present in both.

A refined peak set was generated by calling peaks using MACS2 (CallPeaks, Signac^63^), grouping cells by perturbation condition and cell type, and applying the same peak width and blacklist filters described above. For ATAC-based dimensionality reduction, data were normalized using TF-IDF, top features selected (FindTopFeatures, min.cutoff = 5), and SVD performed (RunSVD). A weighted nearest neighbor (WNN) joint embedding was computed using FindMultiModalNeighbors and visualized using RunUMAP.

Differentially accessible peaks between YBX1 knock-down and unperturbed cells were identified per cell type using FindMarkers with a logistic regression test and nCount_peaks as a latent variable. Differentially accessible peaks were annotated using GREAT (rGREAT R package, genome = hg38^64^) to identify associated genes and distance to the nearest TSS. Gene ontology terms enriched among differentially accessible peak sets were identified using the enrichR R package (v3.2).

A gene activity matrix was created using the Signac GeneActivity function applied to the MACS2 peak set (extend.upstream = 1,000 bp, extend.downstream = 0, biotypes = “protein_coding”), normalized using LogNormalize, and scaled. PFC signature chromatin accessibility per cell was calculated using the module activity scoring approach^24^ applied to the gene activity matrix, with YBX1 excluded from PFC signature gene lists prior to scoring. Chromatin accessibility of individual PFC screen target genes was evaluated analogously using a gene activity matrix computed for all 35 PFC screen target genes.

## Supplementary Table Legends

**Supplementary Table 1.** PFC and V1 marker gene lists extracted from bulk transcriptomic profiling of the developing human cortex (Miller et al., 2014). Table shows gene name, PFC vs V1 marker designation, known links to cortical development, functional annotations, and relevant citations.

**Supplementary Table 2**. Gene list for all 21 area modules generated by our meta-module analysis strategy (Nano et al., 2025).

**Supplementary Table 3.** Relative activity of area modules across developmental stages and cell types in the developing human PFC versus V1 (Nano et al., 2025). Table shows, for each area module, the percent change in module activity in PFC versus V1 samples across combinations of cell type and developmental stage (early (GW 6–17, n = 8), mid (GW 18–20, n = 9), and late (GW 21–26, n = 6)). Only statistically significant changes are shown (p < 0.05, two-sided Wilcoxon test). Modules with greater than 25% increase in activity in the PFC across any combination were designated as PFC signatures.

**Supplementary Table 4.** Gene ontology terms enriched in area modules. Table shows terms enriched in each area module following analysis surveying signaling pathway databases (WikiPathway 2021 Human, KEGG 2021 Human, Elsevier Pathway Collection), transcriptional regulatory collections (ChEA 2016, ENCODE and ChEA Consensus TFs from ChIP, TF Perturbations Followed by Expression, TRRUST Transcription Factors 2019), and gene ontology sets (GO Biological Process 2021, GO Molecular Function 2021, GO Cellular Component 2021). The number of genes in each term (total term size) and the percent and number of term genes included in the indicated module are shown, as well as the adjusted p-value of term enrichment and a list of term genes in the module. All values calculated using the enrichR R package (v3.2).

**Supplementary Table 5.** Annotations for the 18 PFC signatures. Table shows the broad biological process, functional annotation, and cell type- and developmental stage- specificity assigned to each PFC signature early (GW 6–17), mid (GW 18–20), and late (GW 21–26).

**Supplementary Table 6.** PFC screen target genes selected for pooled CRISPR activation screening. Table shows, for each of the 35 PFC screen target genes: gene name; broad biological process category; known links to cortical development (Yes / None/Limited); functional annotation; relevant citations (DOI); the number of stage × cell type contexts in which the gene was identified as a PFC-enriched marker; the cell types in which PFC enrichment was observed (RG, radial glia; IPC, intermediate progenitor cell; Neuron); and the developmental stages in which PFC enrichment was observed (Early, GW 14–17; Mid, GW 18–20; Late, GW 22–25). Marker contexts were identified from a single-cell RNA-seq profile of the developing human PFC and V^14^ as described in Methods.

**Supplementary Table 7.** Library of PFC-targeting gRNAs used for pooled CRISPR activation screening. gRNA sequences were selected from the CRISPRa v2 collection (Horlbeck et al., 2016). For each of the 35 PFC screen target genes, the top 7 sgRNA sequences ranked by predicted activity were included. For target genes with two transcription start sites separated by approximately 1 kb, two independent sets of 7 sgRNAs were designed, one against each TSS, yielding 294 targeting gRNAs in total. Table shows gRNA ID (denoting target gene, strand orientation, genomic coordinates, and TSS set designation), target gene, protospacer sequence, and oligo sequence used for library cloning.

**Supplementary Table 8.** Library of non-targeting (NT) control gRNAs used for pooled CRISPR activation screening. NT gRNAs were selected from the h5 (Gene Expression) sublibrary of the hCRISPRav2 non-targeting collection (Horlbeck et al., 2016) and included at 21-fold greater abundance relative to any individual targeting gRNA, yielding 21 NT gRNAs in total. Table shows gRNA ID, gRNA protospacer sequence, and oligo sequence used for library construction.

**Supplementary Table 9.** Results of context-aware, single-cell gRNA detection in the HPCT pooled CRISPR activation screen (see SFig. 2D). Table shows valid targeting gRNAs for each of the 35 PFC screen target genes.

**Supplementary Table 10.** Cluster markers for single-cell transcriptomic profiles of the HPCT pooled CRISPR activation screen (see Fig. 2F). Markers were calculated using the one-sided Wilcoxon rank sum test and used to assign cell type identities to clusters. Table shows cluster, cluster marker gene name, average log2 fold change, the percentage of cells within the cluster expressing the gene (pct.1), the percentage of cells outside the cluster expressing the gene (pct.2), and adjusted p-value. Positive log-fold change values indicate that the feature is more highly expressed in the cluster of interest.

**Supplementary Table 11.** Table of metadata for all cells in the HPCT pooled CRISPR activation screen (see Fig. 2). Table includes cell barcode, sample of origin, gestational week, annotated cell type, perturbation status for each of the PFC screen target genes (present/absent, as determined by the context-aware gRNA detection pipeline), and overall perturbation classification (unperturbed, perturbed).

**Supplementary Table 12.** Percent change in developmental meta-module (Nano et al., 2025) in singly perturbed versus unperturbed cells in the HPCT pooled CRISPR activation screen (see Fig. 2H). In each of the indicated cell types, the percent change induced by single PFC screen target activation in average activity of each meta-module was calculated relative to unperturbed cells. Table shows cell type, screen target gene, PFC signature, and percent change in activity. Only comparisons with p < 0.05 are shown.

**Supplementary Table 13.** Percent change in PFC signature activity in singly perturbed versus unperturbed cells in the HPCT pooled CRISPR activation screen (see Fig. 2I, SFig. 3D). For each PFC screen target gene, the percent change in average activity of each PFC signature was calculated relative to unperturbed cells, within the indicated cell type or developmental stage. Table shows query population of cells, screen target gene, PFC signature, and percent change in activity. Only comparisons with p < 0.05 are shown. PFC screen target genes were excluded from their respective gene program gene lists prior to scoring.

**Supplementary Table 14.** Spearman correlations between PFC screen target gene expression and PFC signature activity in the HPCT pooled CRISPR activation screen (see Fig. 2K, 2L). Correlations were calculated within cells that were either unperturbed or singly perturbed for the indicated screen target, across all cell types. Table shows screen target gene, PFC signature, and Spearman’s rank correlation coefficient (ρ). Only statistically significant correlations are shown (p < 0.05).

**Supplementary Table 15.** Results of context-aware, single-cell gRNA detection in the hCO pooled CRISPR activation screen (see SFig. 4A, 4B). Table shows valid targeting gRNAs for each of the 35 PFC screen target genes.

**Supplementary Table 16.** Cluster markers for single-cell transcriptomic profiles of the hCO pooled CRISPR activation screen (see Fig. 3B). Markers were calculated using the one-sided Wilcoxon rank sum test and used to assign cell type identities to clusters. Table shows cluster, cluster marker gene name, average log2 fold change, the percentage of cells within the cluster expressing the gene (pct.1), the percentage of cells outside the cluster expressing the gene (pct.2), and adjusted p-value. Positive log-fold change values indicate that the feature is more highly expressed in the cluster of interest.

**Supplementary Table 17.** Table of metadata for all cells in the hCO pooled CRISPR activation screen (see Fig. 3). Table includes cell barcode, sample of origin, gestational week, annotated cell type, perturbation status for each of the PFC screen target genes (present/absent, as determined by the context-aware gRNA detection pipeline), and overall perturbation classification (unperturbed, perturbed).

**Supplementary Table 18.** Percent change in developmental meta-module activity (Nano et al., 2025) in singly perturbed versus unperturbed cells in the hCO pooled CRISPR activation screen (see Fig. 3E, SFig. 4F). In each of the indicated cell types, the percent change induced by single PFC screen target activation in average activity of each meta-module was calculated relative to unperturbed cells. Table shows cell type, screen target gene, meta-module, and percent change in activity. Only comparisons with p < 0.05 are shown.

**Supplementary Table 19.** Percent change in PFC signature activity in singly perturbed versus unperturbed cells in the hCO pooled CRISPR activation screen (see Fig. 3E, SFig. 4G). For each PFC screen target gene, the percent change in average activity of each PFC signature was calculated relative to unperturbed cells, within the indicated cell type. Table shows cell type, screen target gene, PFC signature, and percent change in activity. Only comparisons with p < 0.05 are shown. PFC screen target genes were excluded from their respective gene program gene lists prior to scoring.

**Supplementary Table 20.** Spearman correlations between PFC screen target gene expression and PFC signature activity in the hCO pooled CRISPR activation screen (see Fig. 3I, 3J, SFig. 4H). Correlations were calculated within cells that were either unperturbed or singly perturbed for the indicated screen target, across all cell types. Table shows screen target gene, PFC signature, Spearman’s rank correlation coefficient (ρ), and adjusted p-value. Only statistically significant correlations are shown (p < 0.05).

**Supplementary Table 21.** Cluster markers for single-cell transcriptomic profiles of the YBX1 knock-down hCO experiment (see Fig. 4B). Markers were calculated using the one-sided Wilcoxon rank sum test and used to assign cell type identities to clusters. Table shows cluster, cluster marker gene name, average log2 fold change, the percentage of cells within the cluster expressing the gene (pct.1), the percentage of cells outside the cluster expressing the gene (pct.2), and adjusted p-value. Positive log-fold change values indicate that the feature is more highly expressed in the cluster of interest.

**Supplementary Table 22.** Table of metadata for all cells in the YBX1 knock-down hCO experiment (see Fig. 4).

**Supplementary Table 23.** Percent change in developmental meta-module activity (Nano et al., 2025) in YBX1-KD versus unperturbed cells in the YBX1 knock-down hCO experiment (see Fig. 4D). In each of the indicated cell types, the percent change in average activity of each meta-module was calculated in YBX1-KD cells relative to unperturbed cells from the same organoids. Table shows cell type, meta-module, and percent change in activity. Only comparisons with p < 0.05 are shown. YBX1 was excluded from all gene program gene lists prior to scoring.

**Supplementary Table 24.** Percent change in PFC signature gene expression in YBX1-KD versus unperturbed cells in the YBX1 knock-down hCO experiment (see Fig. 4E-F). For each cell line and cell type combination, the percent change in average gene expression of each PFC signature was calculated in YBX1-KD cells relative to unperturbed cells from the same organoids. Table shows cell line, cell type, PFC signature, and percent change in gene expression. Only comparisons with p < 0.05 are shown. YBX1 was excluded from all PFC signature gene lists prior to scoring.

**Supplementary Table 25.** Differentially accessible chromatin regions between YBX1-KD and unperturbed cells in the YBX1 knock-down hCO experiment, identified per cell type (see Fig. 4G, SFig. 5E). Differentially accessible peaks were identified using the FindMarkers function (Signac; logistic regression test, p < 0.05) with nCount_peaks as a latent variable to correct for sequencing depth. Table shows cell type, peak coordinates, average log2 fold change, the percentage of YBX1-KD cells with accessibility at the peak (pct.1), the percentage of unperturbed cells with accessibility at the peak (pct.2), adjusted p-value, and GREAT-annotated associated genes and distance to nearest TSS. Positive log-fold change values indicate greater accessibility in YBX1-KD cells.

**Supplementary Table 26.** Percent change in PFC signature chromatin accessibility in YBX1-KD versus unperturbed cells in the YBX1 knock-down hCO experiment (see Fig. 4I). For each cell line and cell type, the percent change in average activity of each PFC signature chromatin accessibility was calculated in YBX1-KD cells relative to unperturbed cells from the same organoids. Table shows cell line and cell type, PFC signature, and percent change in activity. Only comparisons with p < 0.05 are shown. YBX1 was excluded from all PFC signature gene lists prior to scoring.

**Supplementary Table 27.** Percent change in PFC screen target gene expression and promoter/gene body chromatin accessibility in YBX1-KD versus unperturbed cells in the YBX1 knock-down hCO experiment (see Fig. 5A, SFig. 6B). For each PFC screen target gene, the percent change in average gene expression and average chromatin accessibility of the gene’s promoter and gene body regions was calculated in YBX1-KD cells relative to unperturbed cells, within each cell type and cell line. Table shows cell line, cell type, PFC screen target gene, percent change in gene expression, and percent change in chromatin accessibility relative to unperturbed cells. Only statistically significant comparisons are shown (p < 0.05, two-sided Wilcoxon rank sum test).

**Supplementary Table 28.** Differentially expressed genes between singly perturbed and unperturbed cells in HPCT deep layer excitatory neurons (see Fig. 5B, 6F, SFig. 6C–H). Genes upregulated or downregulated in cells singly perturbed for YBX1, YBX1-driven PFC screen targets (SOX4, NEUROD6, GTF3A, SMARCE1, ZNF580, MXD4), or YBX1-independent PFC screen targets (ETV1, RORB, MECP2) versus unperturbed cells were calculated using the two-sided Wilcoxon rank sum test. Table shows screen target, gene name, average log2 fold change, the percentage of cells within the perturbed population expressing the gene (pct.1), the percentage of unperturbed cells expressing the gene (pct.2), and adjusted p-value. Positive log-fold change values indicate that the feature is more highly expressed in the perturbed population versus unperturbed controls.

---

**SFig. 1.**
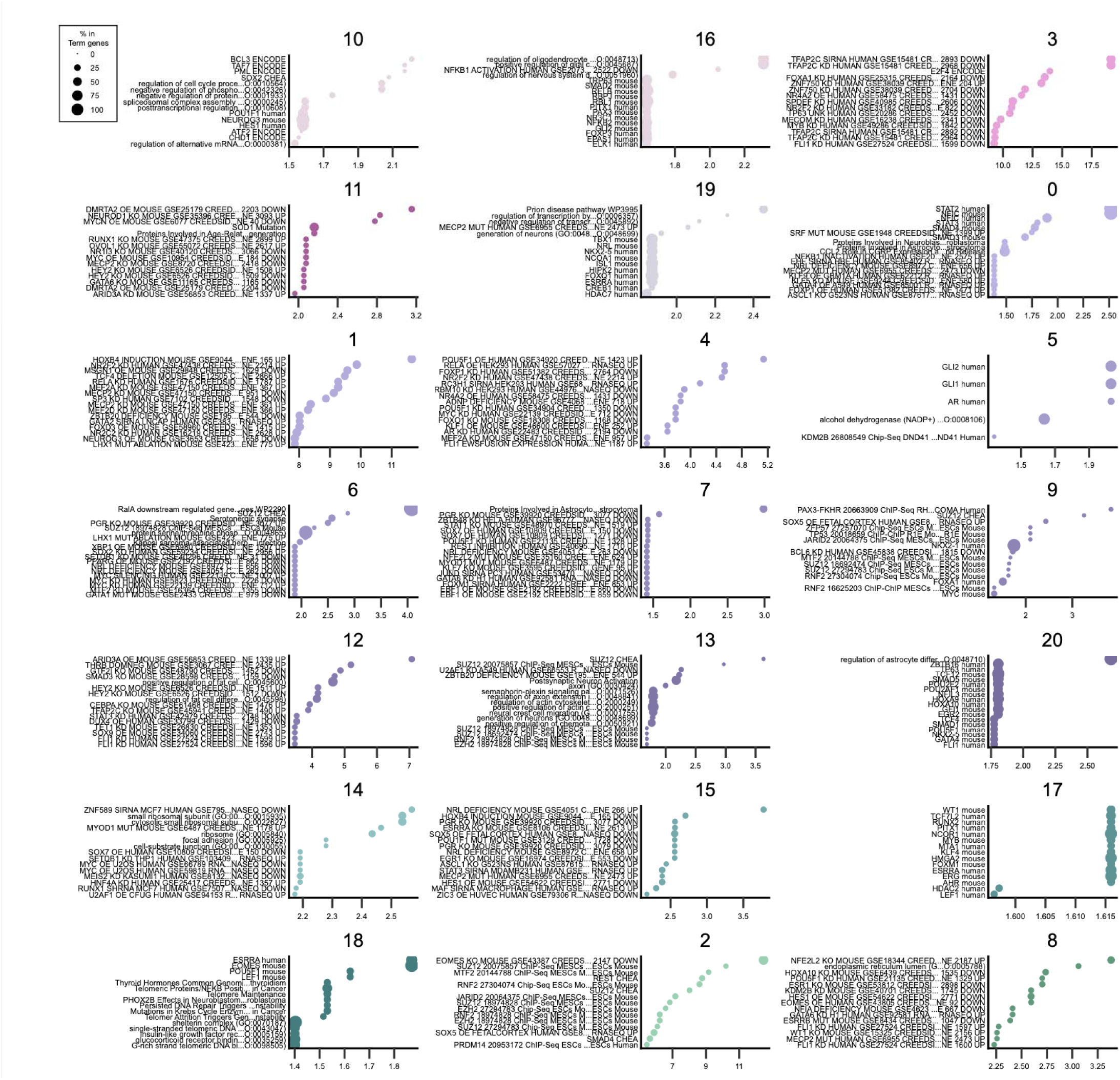
Enriched Gene Ontology Terms in Area Modules. Dot plots displaying gene ontology terms enriched in each of the 21 area modules, including 18 PFC signatures. Dot size reflects the percent of term genes included in the module and dot color reflecting the adjusted p-value.

**SFig. 2.**
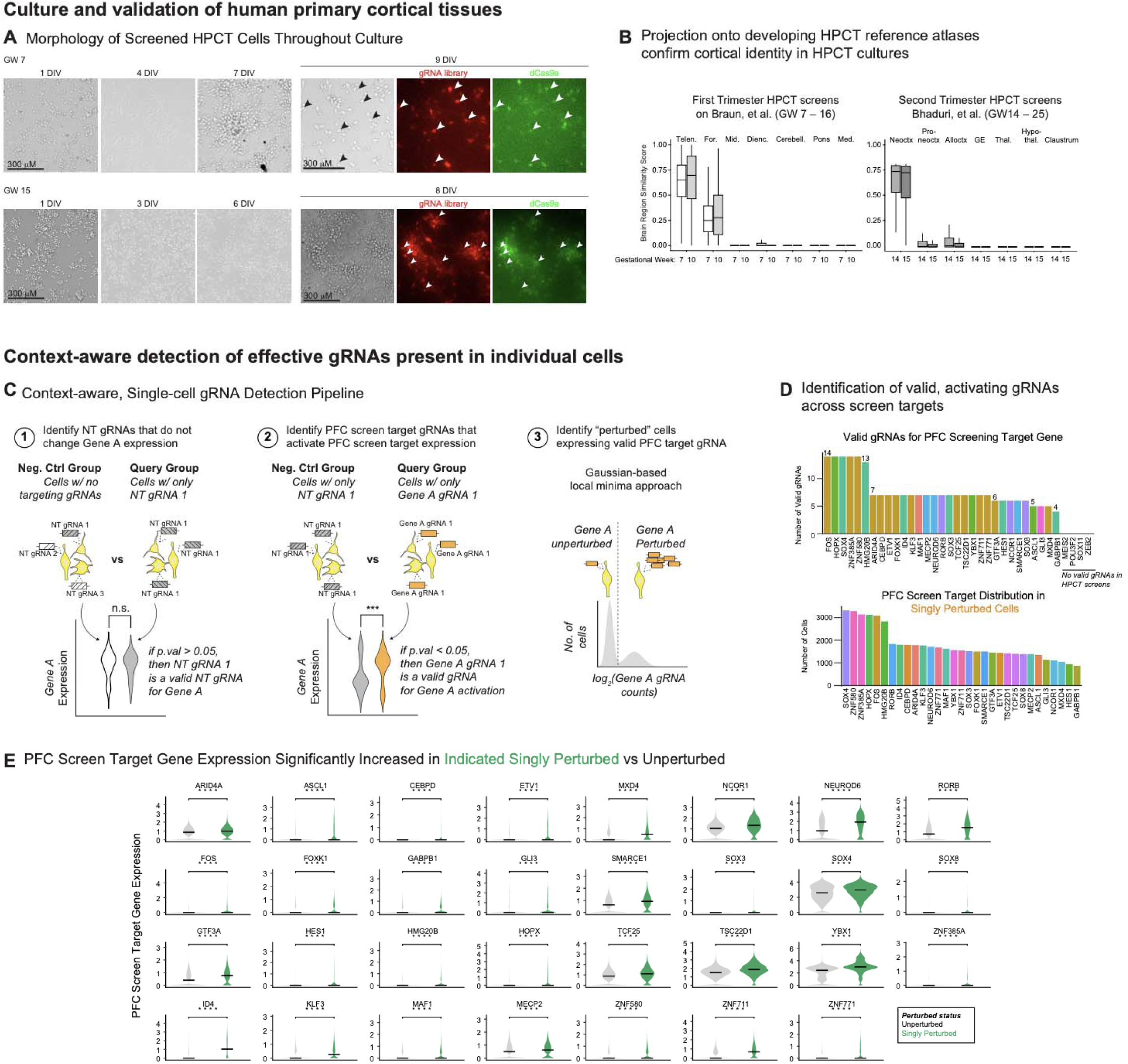
Establishing pooled CRISPR activation screening for areal determinants in HPCT. **A)** Representative live-cell brightfield and fluorescence images of screened HPCT cells throughout the culture period, from 1 days *in vitro* (DIV) to 9 DIV (GW 7; top) or 8 DIV (GW 15; bottom), demonstrating healthy cell morphology and proliferation throughout culture. White arrows, mCherry+; GFP+ cells. Scale bar, 300 μm. **B)** Cortical identity of HPCT cultures was confirmed by projecting screen samples onto HPCT reference datasets: early-stage HPCT screen samples (GW 7 and GW 10) onto a developing HPCT reference atlas (Braun et al.; left) and mid-stage samples (GW 14 and GW 15) onto a second-trimester cortical reference (Bhaduri et al., 2021; right). Boxplots show the resulting similarity scores between screen samples and each brain region present in reference datasets. Screen samples displayed similarity almost exclusively to the telencephalon, forebrain, and neocortex (two-sided Wilcoxon test). **C)** Schematic of the context-aware, single-cell gRNA detection pipeline. For each PFC screen target, the pipeline identifies: (1) non-targeting gRNAs that do not significantly alter gene’s activity (valid NT gRNAs); (2) targeting gRNAs that, relative to valid NT gRNAs, significantly upregulate target gene expression (valid targeting gRNAs); and (3) cells that possess these valid, targeting gRNAs (perturbed cells). Classification of perturbed cells utilized a Gaussian-based local minima approach (Replogle et al., 2020), identifying the cells that met a specific threshold of valid targeting gRNA counts. **D)** Bar charts summarizing the results of context-aware, single-cell gRNA detection. Top, the number of valid, activating gRNAs per PFC screen target. Bottom, the number of cells classified as singly perturbed per PFC screen target. Valid, targeting gRNAs were not detected for MEIS2, POU3F2, SOX11, ZEB2. **E)** Violin plots showing PFC screen target gene expression in unperturbed versus singly perturbed cells for each of the 31 effectively screened target genes, confirming significant activation of each target gene in corresponding singly perturbed cells (****, p-value < 0.0001, two-sided Wilcoxon test).

**SFig. 3.**
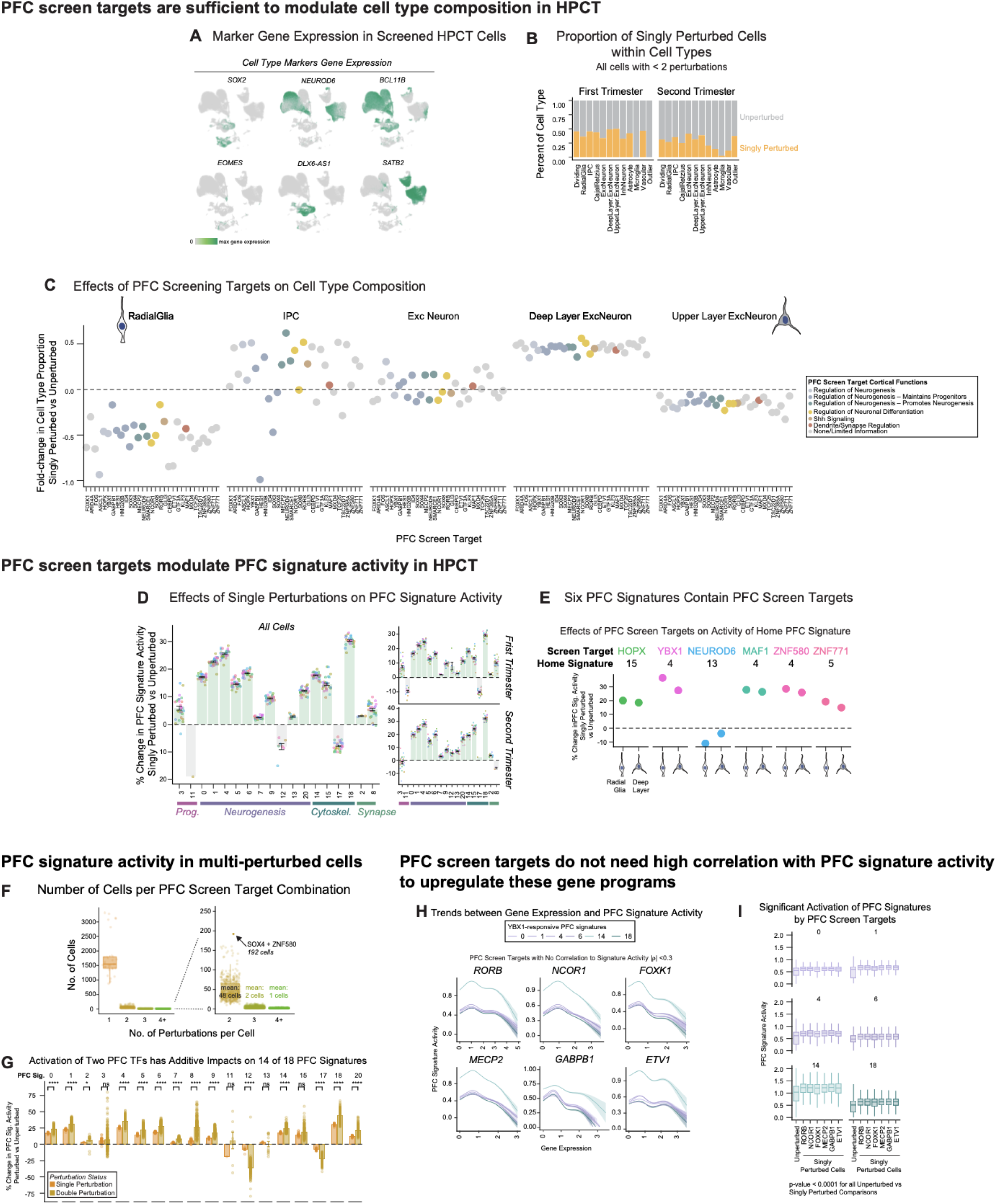
Activation of PFC screen target genes changes cell fate and PFC signatures in HPCT. **A)** UMAPs displaying marker gene expression in screened HPCT cells, including canonical markers *SOX2* (radial glia), *EOMES* (IPC), *NEUROD6* (excitatory neurons), *BCL11B* (deep layer neurons), *SATB2* (upper layer neurons), and *DLX6-AS1* (inhibitory neurons). **B)** Bar charts showing the proportion of singly perturbed cells within each cell type, separately for early-stage (GW 7, GW 10) and mid-stage (GW 14, GW 15) HPCT screens. Perturbed cells are present across all major cortical neuronal cell types. **C)** Dot plots showing the effects of individual PFC screen targets (x-axis) on cell type composition across the glutamatergic lineage. Data presented as fold-change in cell type proportion in singly perturbed versus unperturbed cells. Dots are colored by known cortical functions of each PFC screen target. **D)** Bar and error bars showing the average percent change in PFC signature activity in singly perturbed versus unperturbed cells across all 18 PFC signatures. Analyses were conducted separately for (left) all cell types, (top-right) early-stage and (bottom-right) mid-stage HPCT screens. Effects of individual PFC screen targets shown as individual dots. Only statistically significant effects are shown (p < 0.05, two-sided Wilcoxon test). **E)** Six PFC signatures contain PFC screen target genes: HOPX in signature 15; YBX1, MAF1 and ZNF580 in signature 4; NEUROD6 in signature 13; and ZNF771 in signature 5. Dot plots show the effects of each of these targets on their “home” signature in radial glia and deep layer neurons, with all but NEUROD6 sufficient to activate home signature activity. Only statistically significant effects are shown (p < 0.05, two-sided Wilcoxon test). **F)** Number of cells per perturbed population in HPCT screens. Individual dots represent cells harboring one PFC screen target (1 perturbation) or unique combinations of screen targets in the case of multi-perturbed cells (2, 3, or 4+ perturbations). Inset isolates multi-perturbed cells, showing mean cell counts of 48, 2, and 1 cells respectively. The most frequent double-perturbed combination, SOX4 + ZNF580 (192 cells), is highlighted. **G)** Bar and error bars represent the average percent change in PFC signature activity in perturbed cells vs unperturbed cells across all 18 PFC signatures. Single perturbation (light orange) and double perturbation (dark orange) conditions are shown side by side for each signature. Dots show the effects of individual PFC screen targets (single perturbations) or unique target pairs (double perturbations). Activation of two PFC screen targets has additive impacts on all but 4 of the 18 PFC signatures, with signatures 13, 15, and the two signatures related to progenitor expansion (3, 11) as the exceptions. Only statistically significant effects are shown (p < 0.05, two-sided Wilcoxon test). **H-I)** PFC screen targets that are not highly correlated with PFC signature activity can still activate PFC signatures. **H**) Trend lines represent the relationship between PFC screen target gene expression (x-axis) and PFC signature activity (y-axis). Plots are shown for six targets lacking correlation with PFC signature activity (|ρ| < 0.3) and six PFC signatures demonstrating greatest YBX1 sensitivity (line colors). Trend lines represent estimated smoothed conditional means (general additive model), with shaded region indicating 95% confidence interval. **I**) Boxplots showing PFC signature activity in unperturbed cells versus singly perturbed cells for each of the six screen targets shown in (H). Despite lacking strong correlations with PFC signature activity, all six targets significantly activate these YBX1-sensitive PFC signatures (p < 0.0001 for all unperturbed vs singly perturbed comparisons, two-sided Wilcoxon test).

**SFig. 4.**
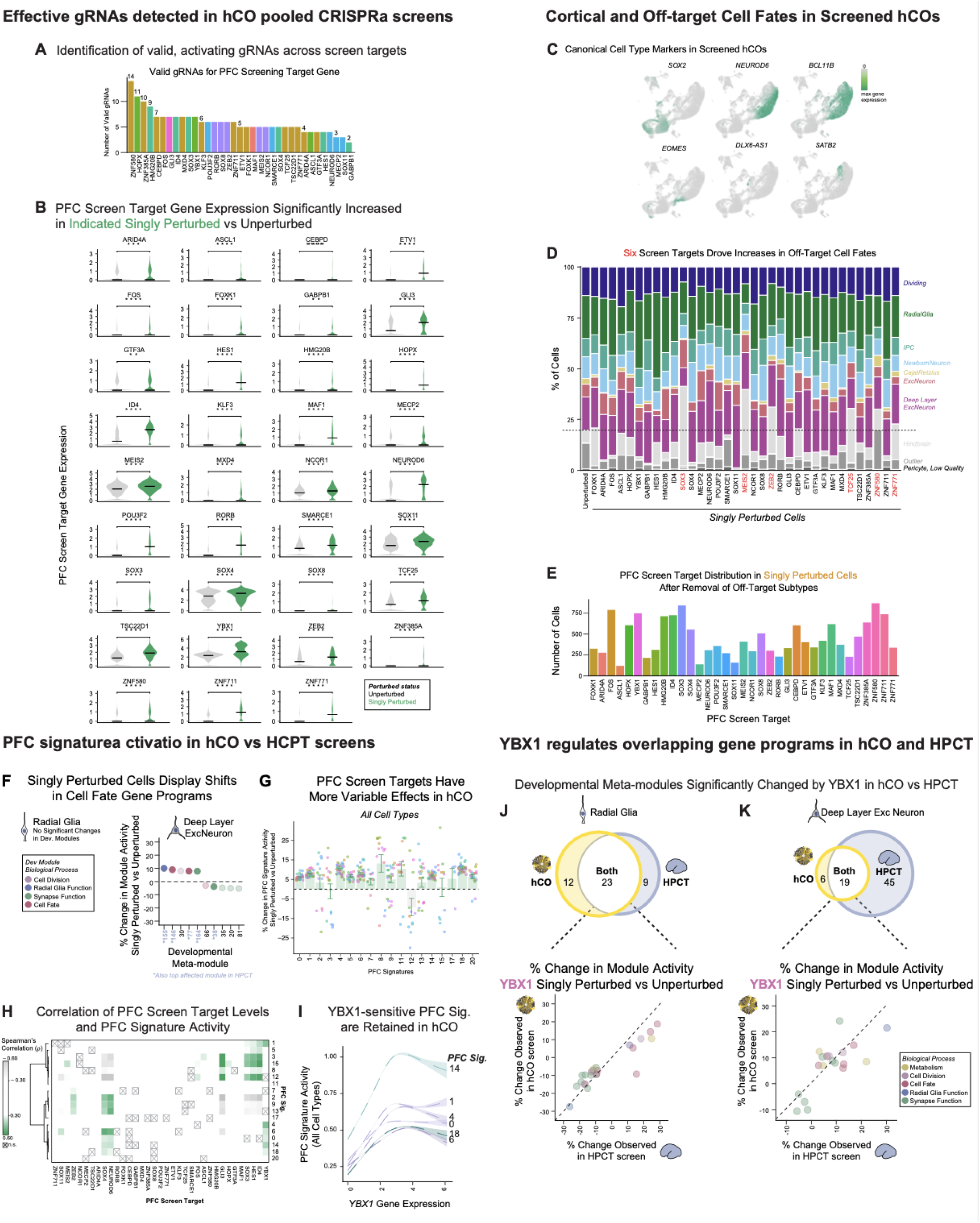
Activation of PFC determinants in human cortical organoids. **A)** Context-aware, single-cell gRNA detection in hCO screens identified a comparable amount of valid, activating gRNAs identified per PFC screen target as in HPCT screens. **B)** Violin plots showing PFC screen target gene expression in unperturbed versus singly perturbed cells for each of the 35 screen targets in hCOs, confirming significant activation of each target gene in corresponding singly perturbed cells (p < 0.05, two-sided Wilcoxon test). **C)** UMAPs displaying canonical cell type marker expression in screened hCO cells, including *SOX2* (radial glia), *EOMES* (IPC), *NEUROD6* (excitatory neurons), *BCL11B* (deep layer neurons), *SATB2* (upper layer neurons), and *DLX6-AS1* (inhibitory neurons). **D)** Stacked bar chart showing the proportion of cortical and off-target cell fates (greyscale) across unperturbed and singly perturbed cells for each PFC screen target. Six screen targets (red) drove increases in hind-brain and other non-cortical cell types relative to unperturbed cells (dashed line). Non-cortical cell types were removed from subsequent analyses. **E)** Bar chart showing the number of singly perturbed cells across PFC screen targets after removal of off-target cell subtypes. **F)** Dot plot showing the percent change in activity of developmental meta-modules (Nano et al., 2025) in singly perturbed versus unperturbed cells. While individual PFC screen targets did not significantly change these cell fate programs in radial glia, screen targets induced modest changes in cell fate programs in deep layer neurons. Developmental meta-modules that were also significantly changed by single perturbations in the HPCT screen are indicated with an asterisk. Only statistically significant effects are shown (p < 0.05, two-sided Wilcoxon test). **G)** Bar and error bars showing the average percent change in PFC signature activity in singly perturbed versus unperturbed cells across all 18 PFC signatures, in all hCO cell types. PFC screen targets have more variable effects on PFC signature activity in hCOs compared to HPCT. Effects of individual PFC screen targets shown as individual dots. Only statistically significant effects are shown (p < 0.05, two-sided Wilcoxon test). **H)** Heatmap displaying correlations between PFC screen target expression and PFC signature activity across all hCO cell types. Only statistically significant measurements shown (p < 0.05, Spearman’s rank correlation test). **I)** YBX1-sensitive PFC signatures retain a positive relationship with *YBX1* expression in hCOs, with PFC signature activity plotted against YBX1 gene expression across all cell types. Trend lines represent estimated smoothed conditional means (general additive model), with shaded region indicating 95% confidence interval. **J–K)** Venn diagrams (top) and scatter plots (bottom) comparing the developmental meta-modules significantly changed by YBX1 in hCO versus HPCT. Analyses conducted separately for radial glia (J) and deep layer excitatory neurons (K). Scatter plots show the percent change observed in the hCO screen versus the HPCT screen for each module, colored by biological process.

**SFig. 5.**
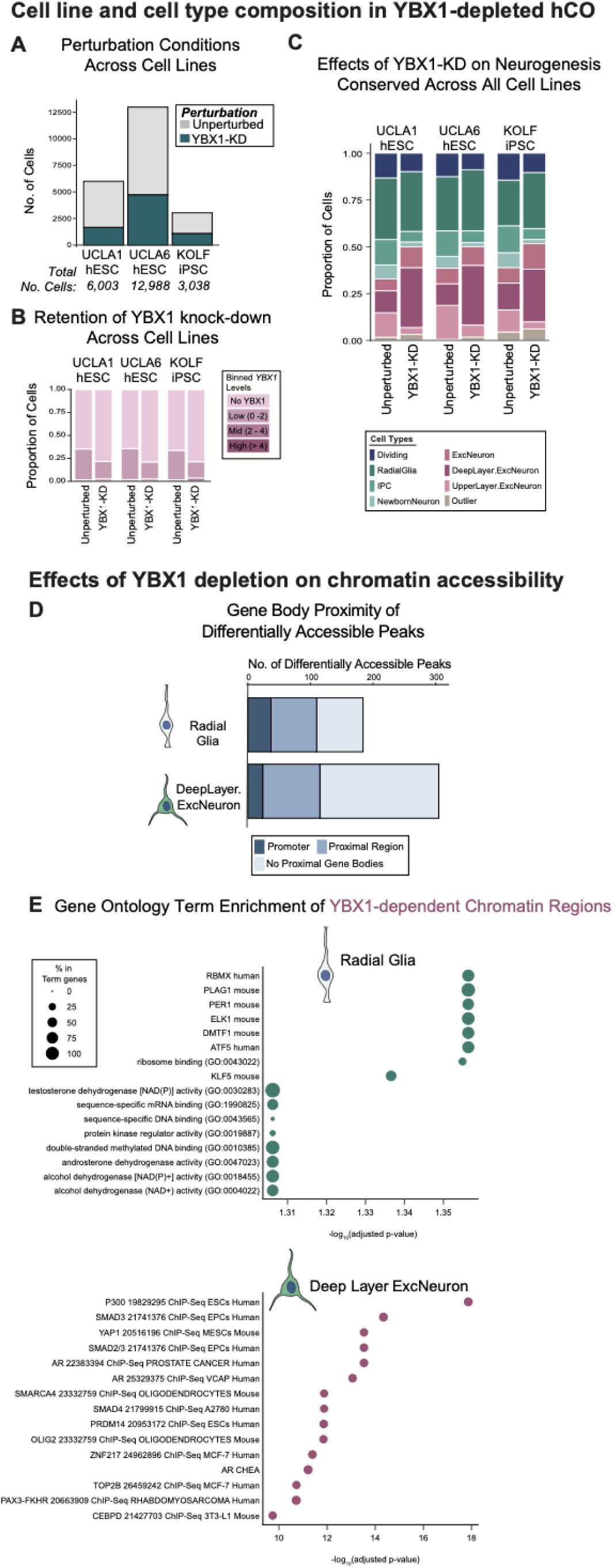
Multi-modal effects of YBX1 depletion in hCOs. **A)** Bar chart shows the number of cells per perturbation condition (unperturbed vs YBX1-KD) across cell lines. **B)** YBX1 depletion is retained consistently across all three cell lines. Bar chart showing the distribution of binned YBX1 expression levels in unperturbed and YBX1-KD cells for each cell line. **C)** Loss of YBX1 consistently expands deep layer subtypes. Stacked bar charts show the effects of YBX1-KD on cell type proportions across all three cell lines. **D)** YBX1 depletion affects a greater number of chromatin regions in deep layer excitatory neurons than in radial glia. Bar chart shows the number of differentially accessible peaks between unperturbed and YBX1-KD cells in radial glia and deep layer excitatory neurons, categorized by proximity to annotated gene bodies (Promoter, Proximal Region, No Proximal Gene Bodies). **E)** Dot plots showing gene ontology term enrichment among the genes associated with YBX1-dependent chromatin regions (regions less accessible in YBX1-KD). Analyses conducted separately for radial glia (top) and deep layer excitatory neurons (bottom). Dot size reflects the percent overlap with term genes and dot color reflects the adjusted p-value.

**SFig. 6.**
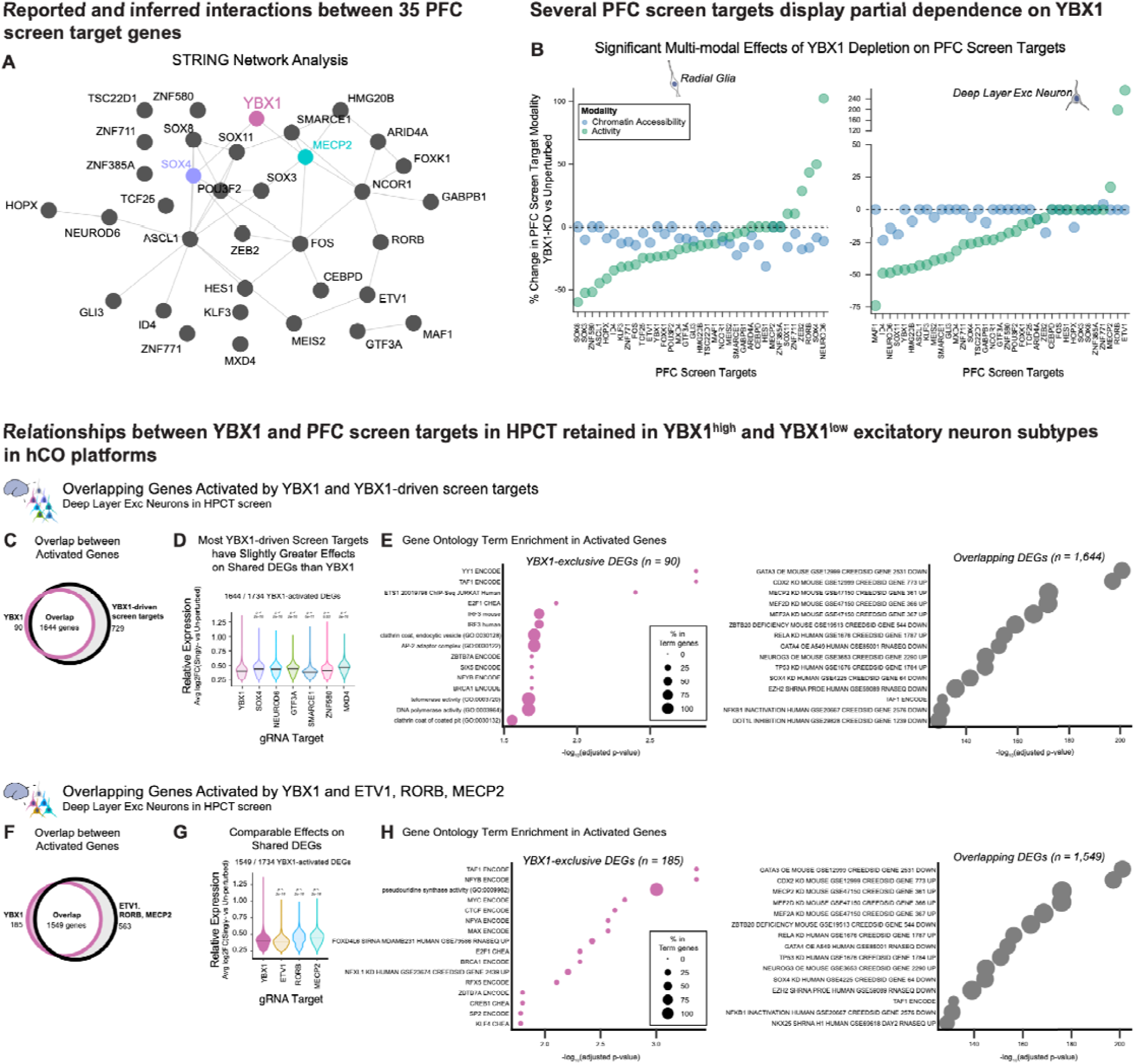
Relationships between YBX1 and PFC screen targets. **A)** STRING network analysis showing reported and inferred interactions among the 35 PFC screen target genes. YBX1 is highlighted, as are its known interactor, MECP2 (Young et al., 2005), and predicted interactor, SOX4. **B)** Several PFC screen targets display significant dependence on YBX1. Dot plots showing the percent change in PFC screen target promoter/gene body accessibility (blue) and gene expression (green) in YBX1-KD versus unperturbed cells. Analyses conducted separately for radial glia (left) and deep layer excitatory neurons (right). Only statistically significant effects are shown (p < 0.05, two-sided Wilcoxon test). **C–E)** Overlap between differentially expressed genes (DEGs) activated by YBX1 and the YBX1-driven screen targets (*SOX4*, *GTF3A*, *ZNF580*, *MXD4*, *NEUROD6*, *SMARCE1*) in HPCT deep layer excitatory neurons. Of 1,734 YBX1-activated genes, 1,644 are also activated by at least one YBX1-driven screen target. There were 90 genes exclusively activated by YBX1 (C, Venn diagram). Most YBX1-driven screen targets have slightly greater effects on shared DEGs than YBX1 itself (two-sided Wilcoxon test, all comparisons to YBX1) (D, violin plots). Gene ontology term enrichment of YBX1-exclusive DEGs (n = 90) and overlapping DEGs (n = 1,644) reveals distinct biological processes enriched in each set (E, dot plots). Dot size reflects the percent of term genes included in the gene set. **F–H)** Overlap between differentially expressed genes (DEGs) activated by YBX1 and the YBX1-independent PFC screen targets ETV1, RORB, and MECP2 in HPCT deep layer excitatory neurons. Of 1,734 YBX1-activated DEGs, 1,549 are also activated by at least one YBX1-independent screen target, with 185 YBX1-exclusive DEGs (F, Venn diagram). The YBX1-independent targets activate a comparable set of DEGs to YBX1 (G, violin plots; two-sided Wilcoxon test, all comparisons to YBX1). Gene ontology term enrichment of YBX1-exclusive DEGs (n = 185) and overlapping DEGs (n = 1,549) is shown (H, dot plots). Genes activated by the YBX1-cohort and by YBX1, MECP2, ETV1, and RORB are enriched for virtually the same biological processes, suggesting that demonstrate that functionally redundant cohorts of PFC determinants converge on a shared transcriptional program in deep layer neurons.

